# SVAT: Secure Outsourcing of Variant Annotation and Genotype Aggregation

**DOI:** 10.1101/2021.09.28.462259

**Authors:** Miran Kim, Su Wang, Xiaoqian Jiang, Arif Harmanci

## Abstract

**Background:** Sequencing of thousands of samples provides genetic variants with allele frequencies spanning a very large spectrum and gives invaluable insight for genetic determinants of diseases. Protecting the genetic privacy of participants is challenging as only a few rare variants can easily re-identify an individual among millions. In certain cases, there are policy barriers against sharing genetic data from indigenous populations and stigmatizing conditions.

**Results:** We present SVAT, a method for secure outsourcing of variant annotation and aggregation, which are two basic steps in variant interpretation and detection of causal variants. SVAT uses homomorphic encryption to encrypt the data at the client-side. The data always stays encrypted while it is stored, in-transit, and most importantly while it is analyzed. SVAT makes use of a vectorized data representation to convert annotation and aggregation into efficient vectorized operations in a single framework. Also, SVAT utilizes a secure re-encryption approach so that multiple disparate genotype datasets can be combined for federated aggregation and secure computation of allele frequencies on the aggregated dataset.

**Conclusions:** Overall, SVAT provides a secure, flexible, and practical framework for privacy-aware outsourcing of annotation, filtering, and aggregation of genetic variants. SVAT is publicly available for download from https://github.com/harmancilab/SVAT

## Background

As the cost of next generation sequencing is decreasing, the number of personal genomes and associated personal information is rapidly increasing. Starting with the initial population-wide genotyping projects such as The HapMap Consortium[1], The 1000 Genomes Project[2], Genomics England[3], The Cancer Genome Atlas (TCGA)[4], Trans-omics for precision medicine (TOPMed)[5], The Genotype-Tissue Expression (GTEx) Project[6], and the Precision Medicine Initiative[7], there are now millions millions of genomes that are deposited in research, clinical, or recreational database. These genomes can provide great insight for developing new therapies and drugs for diseases, prenatal genetic testing[8] and more advanced methods for disease risk prediction[9]. In particular, the high prevalence of genetic data in clinical, recreational, and research areas makes interpretation and management of genomic data challenging. Among these, genomic privacy[10–14] has recently become an important facet of genomic data sharing because the genetic variants are shown to be strong re-identifiers even from de-identified genomic datasets. The sequencing of millions of samples provides genetic variants with allele frequencies spanning a very large spectrum[5, 15]. This makes it even harder to provide privacy as only a few rare variants among millions can easily re-identify an individual[16, 17]. These risks may extend to the relatives of the individuals[18]. While there is a strong urge to share the data for curing diseases, privacy issues are generally not coherently addressed[19]. Recent advances in the usage of DNA as incriminating forensic evidence to solve high-profile cases bring many new ethical questions that may cause concerns for these individuals[17, 20]. Although there is a great hype for open data sharing, there are also policy barriers in sharing datasets especially from vulnerable and indigenous populations and datasets related to stigmatizing conditions.

The size of datasets makes it necessary to outsource the analysis and interpretation of genetic variants are often outsourced to 3^rd^ parties such as cloud-based service providers such as AWS, Azure and Google Cloud[21]. While these services have practically unlimited computational power, the data confidentiality is not always guaranteed as the cloud instance security is very challenging[22]. For instance, the recent breaches of AWS and Solarwind demonstrated that the hackers can perform technically advanced attacks against cloud instances[23]. Industrial standards around “encryption-at-rest” and “encryption-in-transit” are insufficient to protect against IT service compromise.

The field of genomic privacy has grown substantially in recent years. Numerous studies have shown that a small number of variants can lead to re-identification attacks[10, 24, 25] and linking attacks[26–28]. One of the major frameworks that have been successfully applied to genomic data analysis is differential privacy[29, 30] (DP). In DP-based approaches, privacy-enabling data release mechanisms are used to share aggregate statistics from the data. Privacy is enabled by addition of noise such as Laplacian or Gaussian noise[31]. DP has been used to release private GWAS statistics (chi-squared statistics and minor allele frequencies[29]). A common criticism for DP models is the sacrifice of data utility (especially for high dimensional data), which makes it hard to apply in large-scale genomics analysis. Currently, the encryption-based approaches represent the most rigorous route to securely share personal genetic information[32]. There are, however, challenges about their practicality[33]. Methods such as homomorphic encryption[34] provide mathematically provable frameworks for processing encrypted data directly without decryption. Recent studies demonstrated that HE is now potentially practical to perform large-scale genomic computations such as GWAS[35] and genotype imputation[36]. Multiparty computation (MPC)[37] is an alternative cryptographically secure approach, which shares the data into multiple non-colluding entities. The entities communicate intermediate statistics (without leaking any information) with one other in the course of analysis. MPC-based systems may not be practical because they rely on large communication between entities[38]. Several studies developed biomedical data analysis frameworks that combine differential privacy and encryption are proposed[39–42]. These studies demonstrate the practicality of the privacy-enabling data sharing and analysis methods.

In this study, we focus on secure outsourcing of two tasks, namely the variant annotation[43] and aggregation[44–49], which are two basic tasks that are performed in genetic variant analysis pipelines. Annotation is the process of assigning genetic mutations the impact on protein-coding genes. This is a very important first task to characterize the impact of a variant on genes. VEP[50] and Annovar[51] (and accompanying webservers) perform variant annotation using gene annotations and variant list as input[52]. The second task is the aggregation of variants, which is the computation of the variant frequency by counting the existence/allele counts of variants over a large number of individuals. In addition to the annotated impact, the allele frequency provides important insight into a variant’s potential role in diseases[48, 53]. It is used extensively in statistical modeling and machine learning models to estimate the functional impact of variants. For example, NIH’s recently deployed allele aggregation tool, ALFA, which is recently deployed, provides variant aggregation and AF estimation service[45]. In addition, ExAC[47] and GnomAD[49] provide similar services. Aggregation is also utilized in genomic beacons that answer queries about the existence of variant alleles in different databases. It is, however, not clear how the large genotype databases are stored and secured behind these tools.

Our approach, named SVAT, makes use of encryption-based techniques to encrypt the variant data to ensure confidentiality against untrusted entities. The data always stays encrypted while it is being analyzed, which ensures that the cloud-service cannot snoop on the data. Also, even if the encrypted data is hacked or leaked there are no concerns about privacy. SVAT makes use of a vectorized data representation to convert annotations and aggregations into numeric operations, which can be very efficiently computed using Homomorphic Encryption operations. SVAT can perform secure and efficient annotation of SNVs and indels. We also present a memory/time optimization for indels where the annotation is not explicitly dependent on the indel length, SVAT assigns the highest impact with very high completeness. For the aggregation of variants from multiple databases, SVAT utilizes a secure proxy re-encryption approach so that the data can be securely encrypted as multiple sources are combined. Overall, SVAT provides a secure, flexible, and practical framework for privacy-aware outsourcing of annotation, filtering, and aggregation of genetic variants.

## Results

We first review the secure annotation and aggregation methods that SVAT implements. We next present the comparison with plaintext annotation/aggregation and computational requirements.

### Overview of Secure Variant Annotation and Aggregation

We present the approaches for annotation (Figure 1a) and aggregation (Figure 1b) tasks.

**Figure 1.**
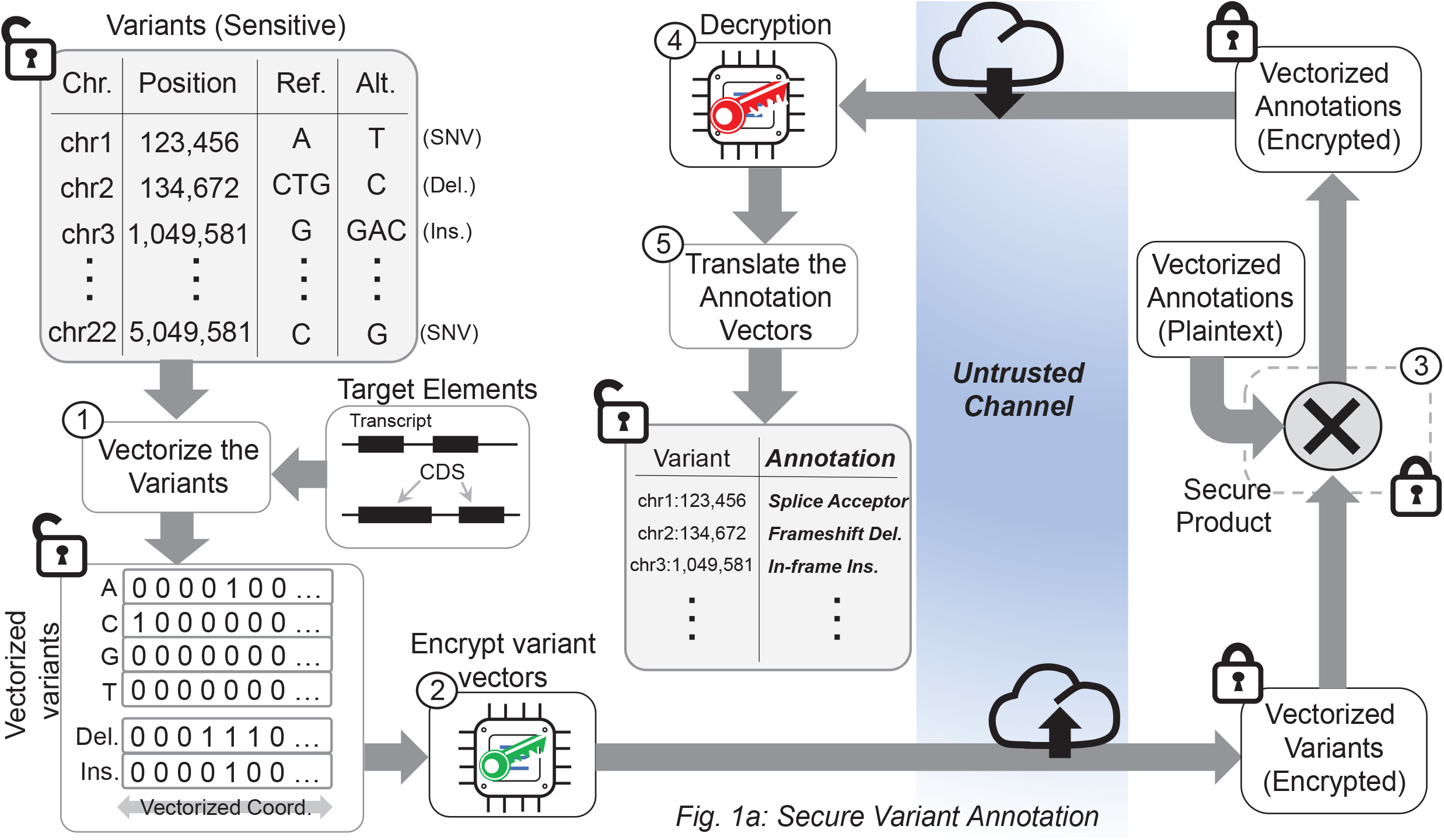
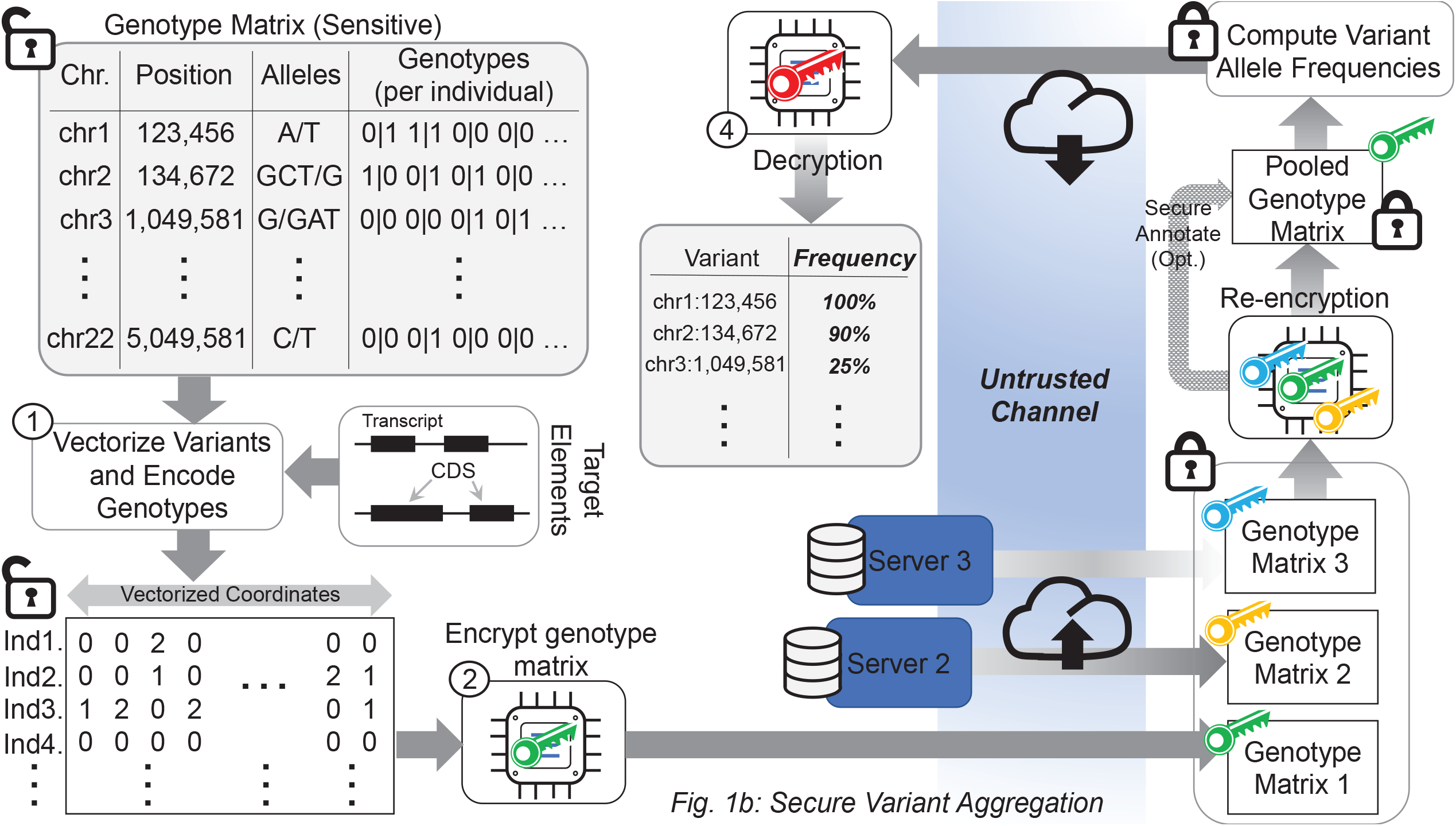
The illustration of annotation, aggregation tasks. (a) The secure annotation task starts by vectorization of the variant loci on the target regions that cover multiple transcripts. The vectorized variant loci signal is encrypted by the researcher’s public key and submitted to the annotation server. The server performs secure multiplication of the variant loci signal and the impact signals and generates the annotated vector that is encrypted. The annotated vector is sent back to the researcher. The researcher decrypts the signal using the private key and translates the annotated variants. (b) The secure aggregation task starts by vectorizing the variant loci. Next, the matrices are encrypted by the owners. The encrypted genotype matrices from multiple databases are stored at the aggregation server. When an aggregation task is requested by a researcher, the server re-encrypts the matrices using re-encryption keys so that it can be decrypted by the researcher’s private key. The re-encrypted matrices are pooled and securely aggregated to compute the allele frequency at each position on the vectorized positions. The resulting frequency array (encrypted) is sent back the researcher and is decrypted by the researcher.

#### Variant Annotation

Figure 1a illustrates the secure variant annotation process. Variant annotation takes the set of target regions (i.e., a BED file), the variant coordinates (e.g., a VCF file), and gene annotations as input and annotates each variant with an impact value, e.g., “splice_acceptor”, or “missense_mutation”. The target regions are used to filter the variants to the regions of interest such as protein-coding sequences or exons. The gene annotations are input as GFF/GTF files and they are central for variant annotation to describe the exact position of the coding sequences so that the untranslated regions (UTRs), start/end codons, and coding frames can be explicitly identified for each gene and transcript. SVAT utilizes GENCODE gene annotations[54] by default. We exclude the transcripts that are annotated as nonsense-mediated decay and incomplete coding sequence because these transcripts may contain incomplete annotations coming from incomplete evidence.

#### Selection of Target Regions

The target regions are used to decrease the amount of data that needs to be submitted between the researcher and the annotation/aggregation server. SVAT utilizes, by default, the protein-coding transcript exons or coding sequences (CDS), which harbor the most consequential phenotype-impacting mutations on the genome (Fig. 2a). In addition, SVAT extends the ends of each exon by a certain length (10 base pairs by default) to include the variants in the introns that may impact splice acceptor/donor motifs. For GENCODE v31, we identified 754,713 protein-coding exons for 83,666 protein-coding transcripts covering 205,443,663 nucleotides. Further, when we exclude the untranslated regions (UTRs), the target regions (with only coding sequences) cover 113,479,849 nucleotides. The exclusion of UTRs is reasonable since UTRs contain variants that are relatively much lower impactful mutations compared to the coding sequences.

**Figure 2.**
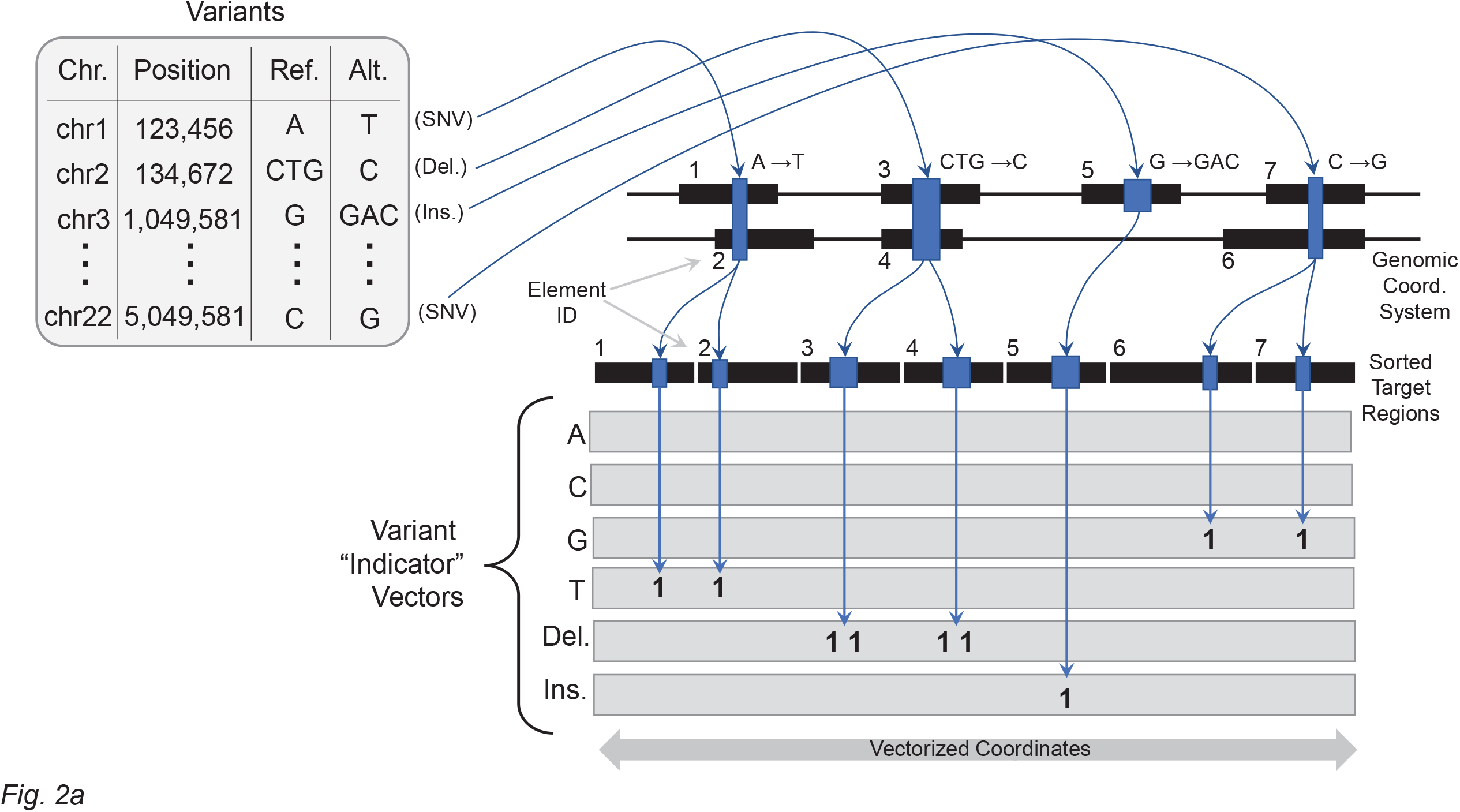
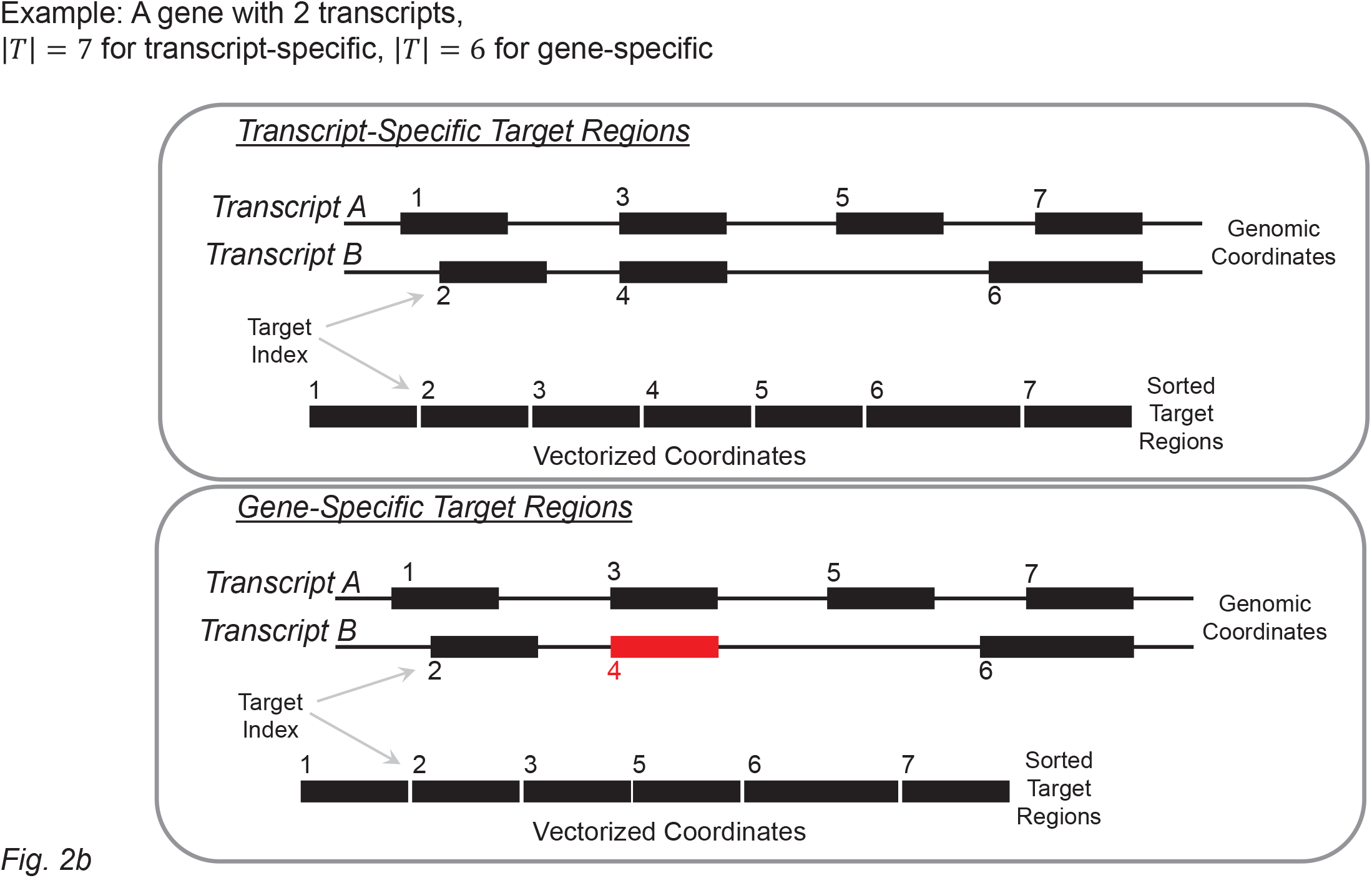
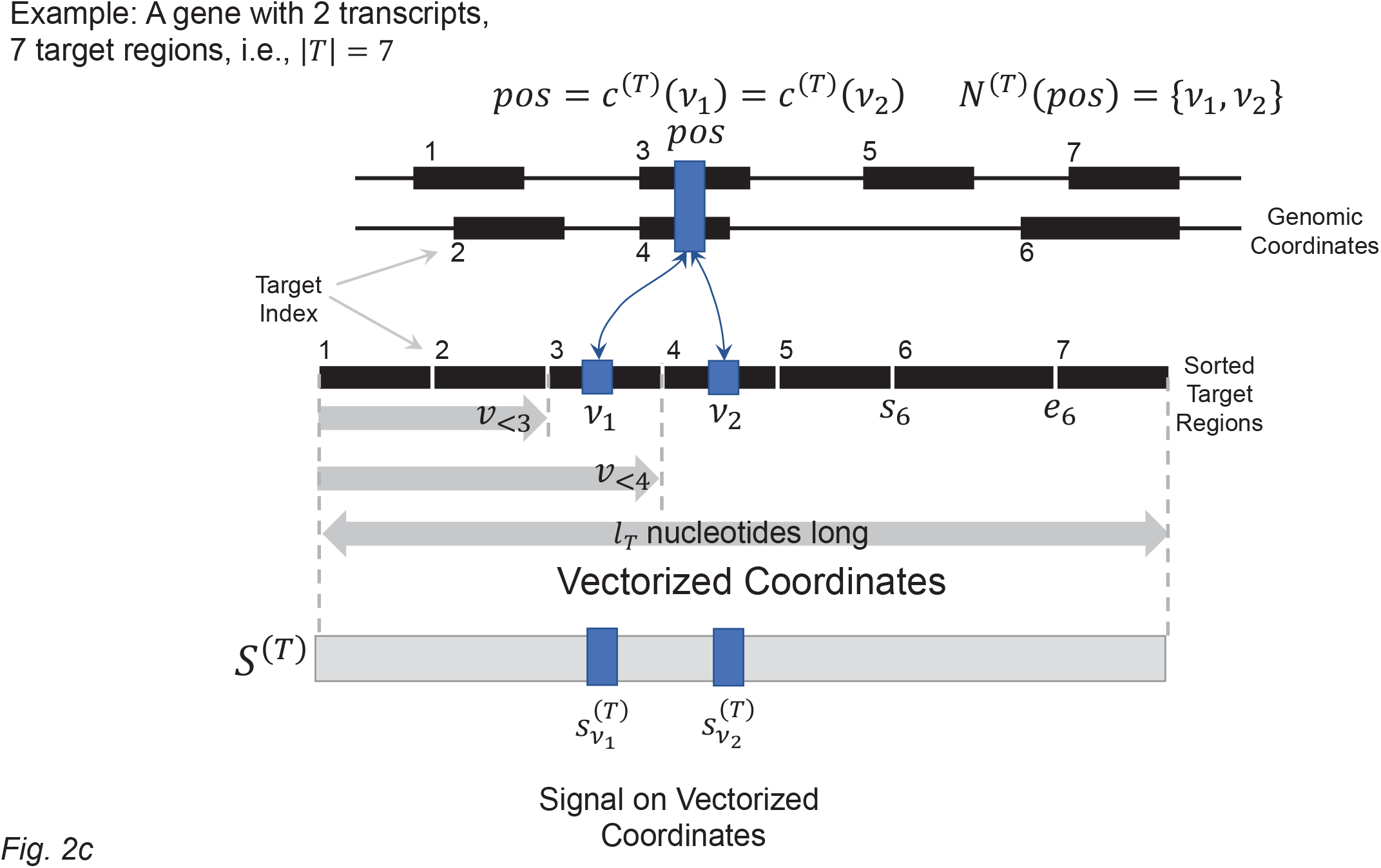
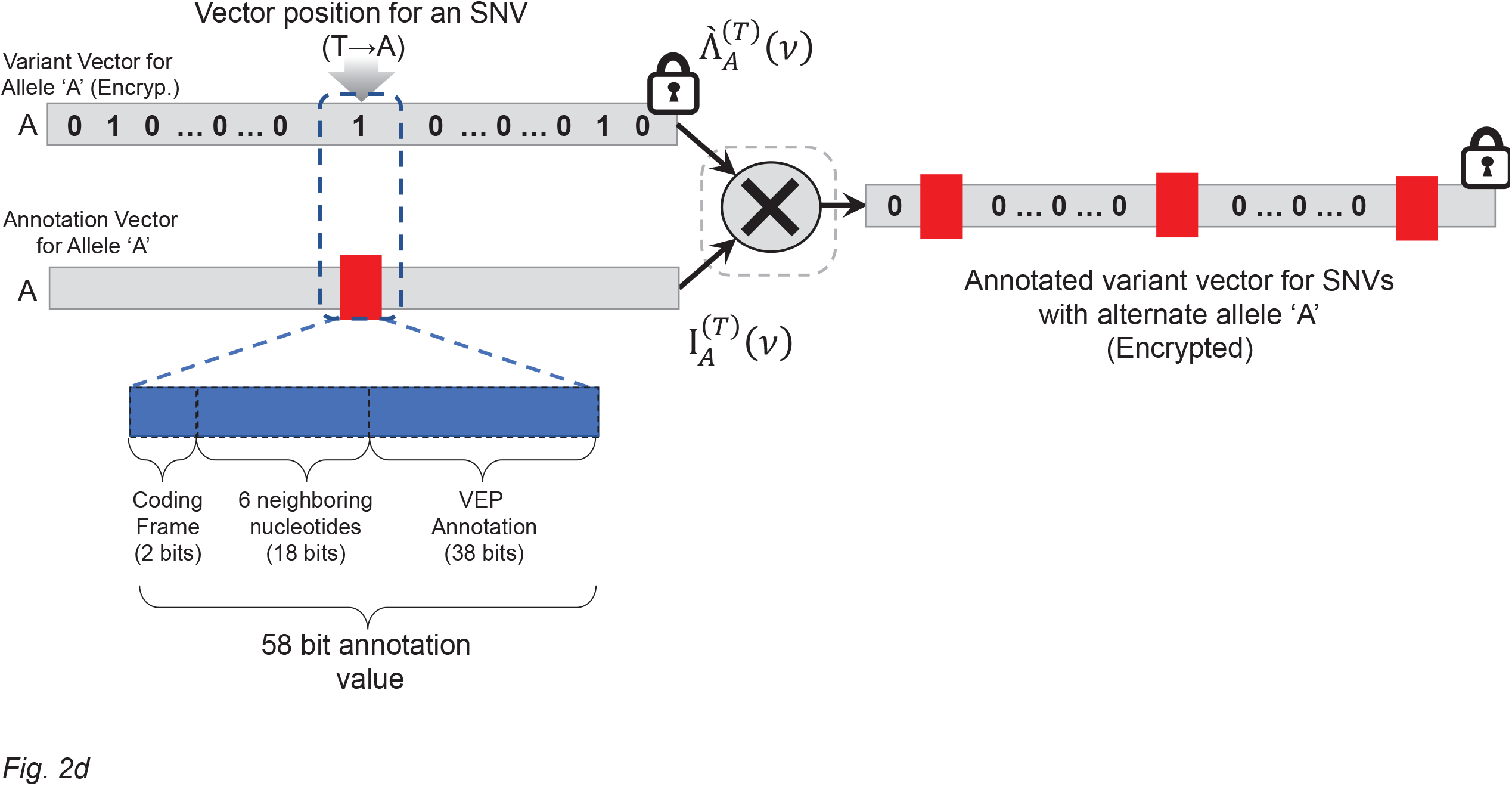
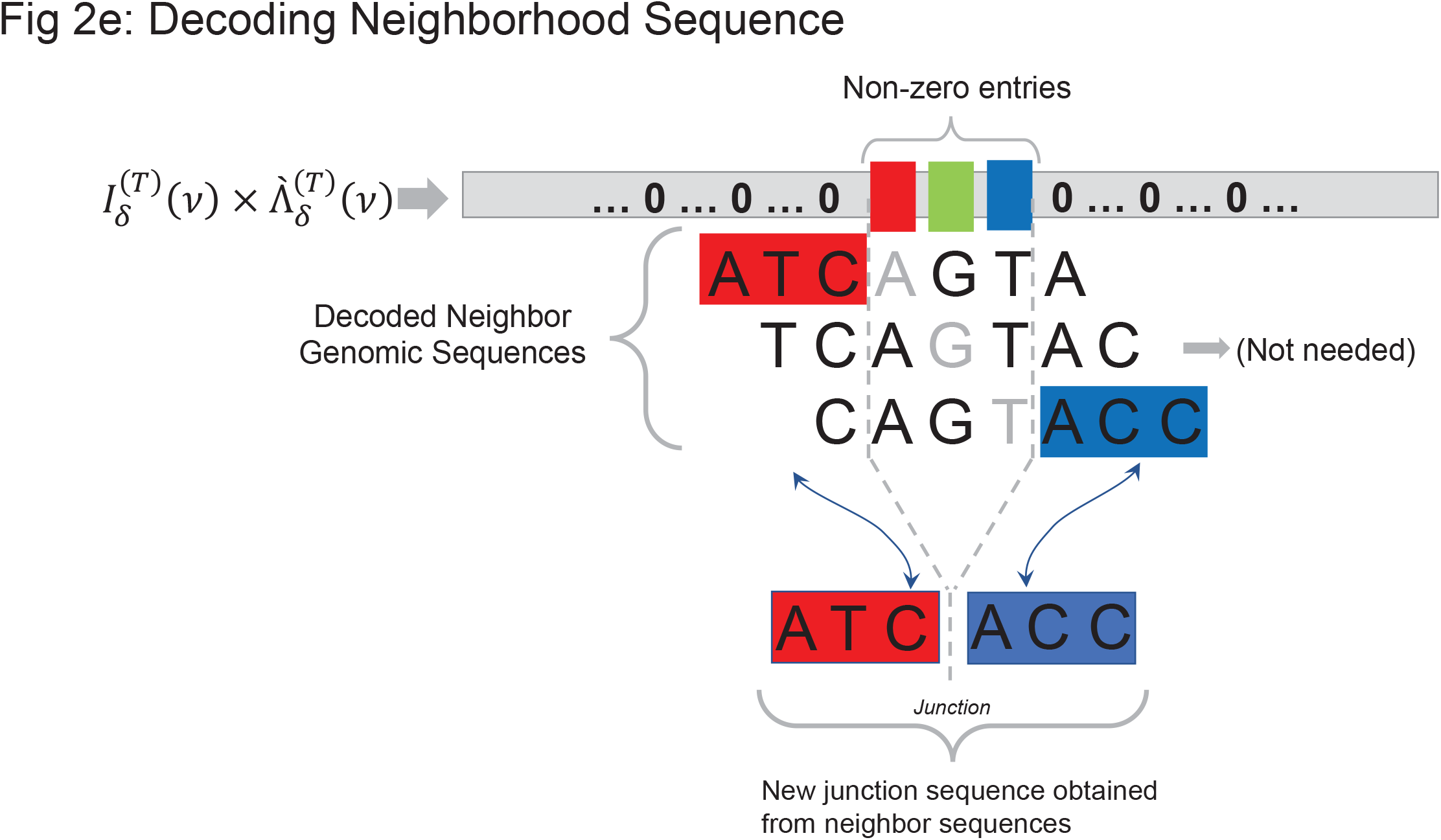
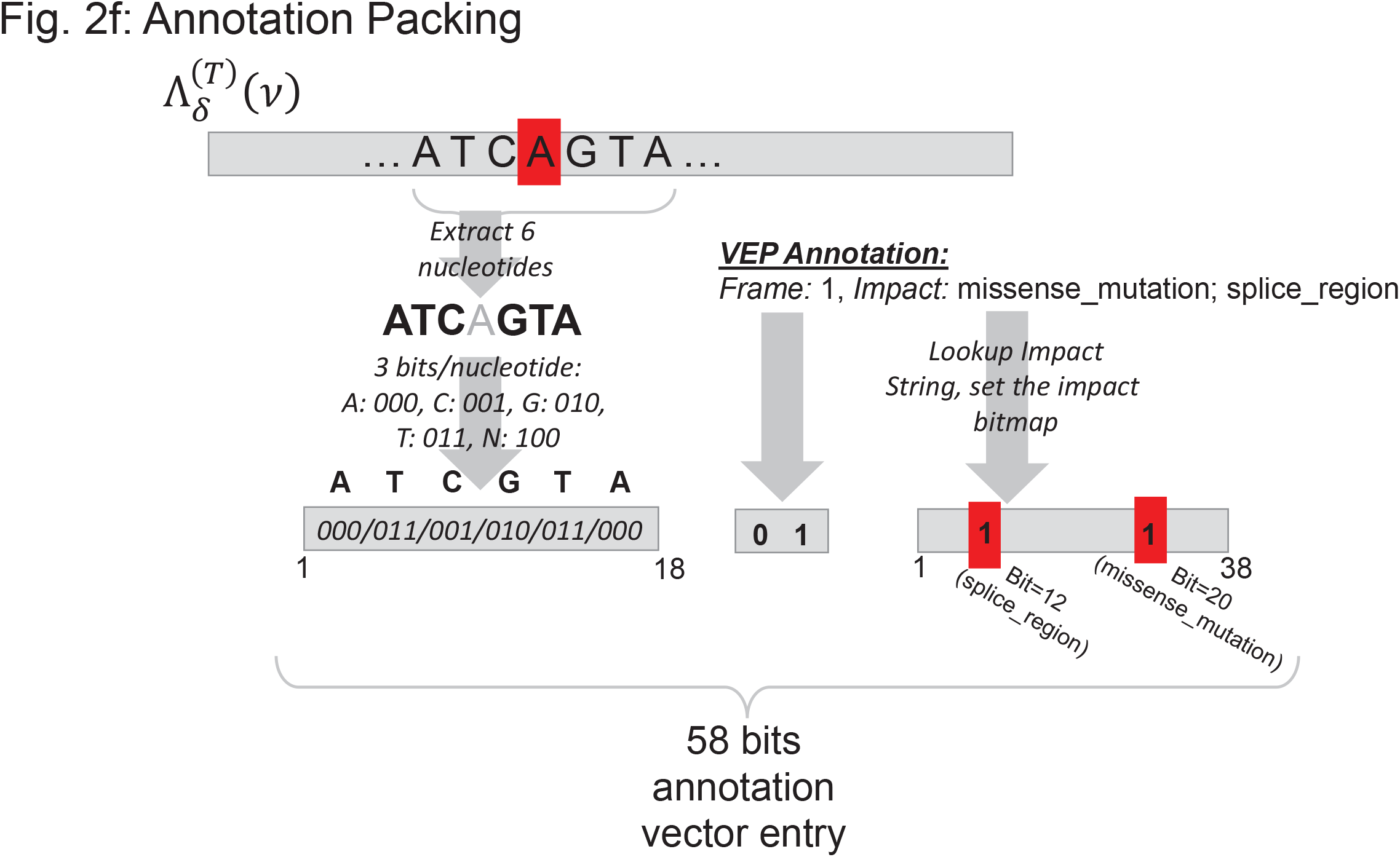
Illustration of the variant loci vectorization. (a) Example of variant locus vectorization over 7 target regions of 2 transcripts. 4 mutations are shown as they are mapped on the vectorized coordinates. Two SNVs overlap with both transcripts, the corresponding positions on targets 1,2 for 1^st^ SNV and targets 6,7 for 2^nd^ SNV are set to “1” for the vector that is corresponding to the alternate alleles of the SNVs. Similarly, 2 base-pair deletion is mapped to the vectorized positions impacted by the deletion and 2 consecutive entries in the vector are set to “1”. For the insertion, the position corresponding to the insertion is set “1”. (b) Example of transcript-specific (Top) and gene-specific (Bottom) target regions. The gene-specific regions do not contain one exon that is redundant in terms of variant annotation. (c) Illustration of the notations. (d) Illustration of the multiplication between the impact vector (58-bit annotation value vector) and the variant locus vector. The multiplication is indicated by the cross and the results vector is shown on the right. Any non-zero entry is illustrated by red rectangles. (e) Example decoding of the junction sequence using the neighborhood sequences for a 3-nucleotide deletion. The junction sequence is formed simply by joining the left and right neighborhood sequences of the first and last nucleotides of the deletion. (f) The 58-bit packing of variant annotation.

These target regions contain all possible alleles for all transcripts. We thus refer to it as “transcript-specific target regions”. We observed that there are substantial number of CDSs in alternatively spliced genes where the frame and start/end coordinates are exactly matching. From a variant annotation point of view, these CDSs provide redundant information because any variant that overlaps one of these CDSs will have exactly the same impact on all the CDSs. To decrease the redundancy, SVAT assesses all the CDSs of each gene and uses the CDSs that are unique when the start/end position and the coding frame are considered. It should be noted that this list excludes the majority of the untranslated regions (UTRs) since the focus is on CDSs (with an end extension of 10 bp by default). This new set of target regions represent the targets in a gene-specific manner: For each gene, all the CDSs at every possible frame are included. Thus, we refer to these target regions as “gene-specific target regions”. The advantage of using gene-specific target regions can decrease the computational requirements. We analyzed the 19718 protein-coding gene annotations in GENCODE v31 and we identified 259,478 exons that cover 48,041,893 nucleotides in the gene-specific target regions. The gene-specific target regions can be used when the focus is on identifying the most impactful annotation of a mutation among multiple transcripts of a gene, which is the most common way that the mutations are summarized among multiple transcripts (Fig. 2b).

As the vectorized loci must represent every position on every transcript, the genomic coordinates will be redundantly represented on the vectorized loci. This is necessary because we would like to report the impact of any variant on overlapping transcripts which share exons.

#### Adversarial Model

The privacy and confidentiality of the genetic variants and genotypes are treated as sensitive information. We assume that the variant annotations are not explicitly sensitive, and therefore the researcher is not an adversarial entity. In this scenario, the researcher’s privacy (the privacy of the participants) needs to be protected against the untrusted annotation server, which we assume does not act maliciously (does not deviate from the protocols and does not collude with other users) but may want to snoop into the contents of the variant and genotype data if they can.

There are two sensitive components for the genetic data: (1) Variant locus and alleles, i.e., chromosome, position and alternate allele, and (2) the genotypes of the variants for each participant. For the variant annotation task, we assume the variant genotypes are not required and only the variant loci and alleles are necessary. This is reasonable since the impact is generally evaluated with respect to only the position and the alternate allele of the variants. We, therefore, will assume the annotation task requires protection of the variant loci and the alternate allele information.

In order to protect the variant loci, it is necessary to protect the locations of the variants, for example, by encrypting the chromosome and position values. Next, the encrypted loci can be annotated using, for example, a private (or secure) set intersection protocol, which may induce a high communication cost between the researcher and the annotation server such that the client and the server must be up and responsive to the communication protocols. SVAT utilizes a framework based on homomorphic encryption (HE), which encodes the variants into a vectorized array, then encrypts the sensitive loci and genotype data once at the researcher’s computer. The data is encrypted while in transfer and while it is being processed at the annotation server. After the annotation is finished, the researcher receives the annotated data, decrypts it and translates the vectorized annotation information. We describe these steps below:

#### Vectorized Representation of the Variant Loci

The vectorization is necessary to protect the variant loci that fall on the target regions. The idea is to enumerate the mutations (i.e., SNVs) (All positions on all transcripts and all alleles) and use the vectorized array for any annotation task. This way, the untrusted entity always receives the encrypted mutational status of all nucleotides (of all alleles) on the target regions regardless of whether there is a mutation or not. Thus, it will not learn anything about the variant loci. The main advantage of the vectorization is that the vector coordinates are not sensitive since they are standardized and do not leak any information (Fig. 2c). Also, as practical homomorphic encryption systems such as Brakerski-Gentry-Vaikuntanathan (BGV)[55], Brakerski/Fan-Vercauteren (BFV)[56, 57], Fully Homomorphic Encryption over the Torus (TFHE)[58], and CKKS[59] support homomorphic operations on encrypted vectors, we only need to use a conventional encryption method of the system and no further elaboration is necessary for homomorphic computation. Thus, the vectorized mutations can be processed using packing and streaming operations to improve performance[33].

We describe the general steps of the vectorization process, which aims at producing a linearly indexed array (i.e., a vector) from the overlapping target regions on the genome (See Methods for details). SVAT first extracts the start/end coordinates of the target regions. Each target region (e.g., a protein-coding exon) is extended (by *l*_*ext*_ base pairs) to include variants that may impact splicing. Next, SVAT sorts the extended target regions with respect to first to start positions and then the element ids (e.g., gene/transcript name). Next, the sorted target regions are “stitched” together (i.e., connect the leftmost end of each region to the rightmost end of the region before this region) in the order that they are sorted (Fig. 1a). After stitching, every position on the target regions can be indexed by one index value on the stitched array, which is the final vectorized array. As we have discussed above, the vectorized array contains as many positions as the total number of nucleotides (which we denote by *l*_*T*_) covered by the target regions. Every position on the vectorized array can be mapped to a unique position on the target regions and vice versa.

Clearly, a position on the genome can map to multiple positions on the vectorized representation since the genomic position may overlap with multiple target regions, i.e., multiple CDSs of a gene. To map a genomic coordinate to the vectorized coordinates, we first identify the target regions that overlap with the genomic coordinate. Next, the positions on the target regions are mapped to the vector coordinates. Since the target regions are sorted with respect to the start position, the searching of genomic coordinates within sorted target regions can be efficiently performed (Fig. 2b).

#### Vectorized Mutation Loci

Given the target regions *T*, SVAT allocates a vector of length *l*_*T*_ (indexed by the vectorized coordinates) and stores the variant locus information on the vector. Given a variant allele *a* (*a E* {*A, C, G, T, δ, L*} the nucleotides of SNV, and deletion and insertion (representing 1-base-pair, i.e., 1-bp, deletion and 1-bp insertion), an array of *l*_*T*_ is allocated, and each array entry whose vectorized position overlaps with a variant is set to 1 (Fig. 2b,c). All other positions are set to 0 (Fig 2d):

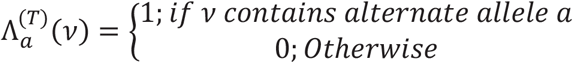

where 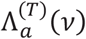 denotes the variant loci array for allele *a*. The above equation simply describes that the position *v* on the mutation loci array is set to 1 if there exists a mutation with alternate allele *a* whose genomic coordinate maps to *v*. For deletions, a separate variant loci array can be generated for different deletion lengths and set the position where the deletion starts as 1 in the array. Alternatively, SVAT makes use of 1-bp deletion arrays such that each deletion is treated as a sequence of 1-bp deletions and 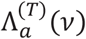 is set to 1 for every position that overlaps with a deletion. This is a more efficient representation because only one variant loci vector is sufficient. For insertions, each variant is described by the position where insertion manifests. The array index that maps to the insertion’s coordinate is set to 1.

#### Encryption of the Vectorized Variant Loci

The researcher (or the data owner) provides the public key for the encryption. The. Based on the encryption parameter setting of an underlying homomorphic encryption system, we denote *l* to be the maximal length of a plaintext vector. Then, an array of length *l*_*T*_ is divided into plaintext vectors of size less than *l*, each of which is encrypted using the public key of the system.

#### Vectorized Annotation Information

Similar to the variant locus array, a vector of length *l*_*T*_ stores the annotations of all mutations on the target regions (Fig. 2d). This is very similar to the mutation loci vector, except that for each position *v*, the array stores a 58-bit entry that packs the annotation information:

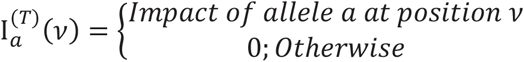

The impact value indicates the impact of the mutation located at vector position *v* with allele *a*. The impact information is retrieved from a variant annotation tool, VEP[50], by default. SVAT packs the impact information and the nucleotide information (Fig. 2e) around the mutation locus. As illustrated in Figure 2f, the packed impact information contains:

1. Coding frame (2 bit): This is the coding frame of a mutation that is a value of out of the set {0,1,2}
2. 6-base pair nucleotide neighborhood (18 bits in total): SVAT uses 3-bits for each nucleotide and extracts an 18-bit value to encode the 6-base pair vicinity of the mutation, which is the one-codon neighborhood of each position.
3. The assigned impact string identifier assigned by VEP (38 bits): A bitmap (38-bits) that describes the impact values of the mutation out of the 38 impact values that VEP assigns to each mutation. The bitmap is generated by setting the bits to 1 for the indices that correspond to the impact strings assigned by VEP. Table 1 shows the impact strings that SVAT uses.

**Table 1:** VEP Impact terms that impact coding gene regions. First column shows the impact term. Each cell with ‘1’ indicates the gene element that is impacted. The full set of impact terms are provided in Supplementary Table 1 in Additional File 1.

| *Term* | *CDS* | *Splice*<br>*Region* | *Splice*<br>*Acceptor* | *Splice*<br>*Donor* | *5'*<br>*UTR* | *3'*<br>*UTR* | *Start*<br>*Codon* | *Stop*<br>*Codon* | *Intron* |
| --- | --- | --- | --- | --- | --- | --- | --- | --- | --- |
| *intron_variant* | 0 | 0 | 0 | 0 | 0 | 0 | 0 | 0 | 1 |
| *3_prime_UTR_variant* | 0 | 0 | 0 | 0 | 0 | 1 | 0 | 0 | 0 |
| *5_prime_UTR_variant* | 0 | 0 | 0 | 0 | 1 | 0 | 0 | 0 | 0 |
| *coding_sequence_variant* | 1 | 0 | 0 | 0 | 0 | 0 | 0 | 0 | 0 |
| *synonymous_variant* | 1 | 0 | 0 | 0 | 0 | 0 | 0 | 0 | 0 |
| *stop_retained_variant* | 0 | 0 | 0 | 0 | 0 | 0 | 0 | 1 | 0 |
| *start_retained_variant* | 0 | 0 | 0 | 0 | 0 | 0 | 1 | 0 | 0 |
| *incomplete_terminal_codon_variant* | 1 | 0 | 0 | 0 | 0 | 0 | 0 | 0 | 0 |
| *splice_region_variant* | 0 | 1 | 0 | 0 | 0 | 0 | 0 | 0 | 0 |
| *protein_altering_variant* | 0 | 0 | 0 | 0 | 0 | 0 | 0 | 0 | 0 |
| *missense_variant* | 1 | 0 | 0 | 0 | 0 | 0 | 0 | 0 | 0 |
| *inframe_deletion* | 1 | 0 | 0 | 0 | 0 | 0 | 0 | 0 | 0 |
| *inframe_insertion* | 1 | 0 | 0 | 0 | 0 | 0 | 0 | 0 | 0 |
| *transcript_amplification* | 0 | 0 | 0 | 0 | 0 | 0 | 0 | 0 | 0 |
| *start_lost* | 1 | 0 | 0 | 0 | 0 | 0 | 1 | 0 | 0 |
| *stop_lost* | 1 | 0 | 0 | 0 | 0 | 0 | 0 | 1 | 0 |
| *frameshift_variant* | 1 | 0 | 0 | 0 | 0 | 0 | 0 | 0 | 0 |
| *stop_gained* | 0 | 0 | 0 | 0 | 0 | 0 | 0 | 1 | 0 |
| *splice_donor_variant* | 0 | 0 | 0 | 1 | 0 | 0 | 0 | 0 | 0 |
| *splice_acceptor_variant* | 0 | 0 | 1 | 0 | 0 | 0 | 0 | 0 | 0 |

In order to generate the vectorized annotation, VEP is run to generate the annotation of the mutations on all nucleotide positions of the target regions and all alleles. For deletions, VEP can be either run for multiple deletion lengths, or SVAT can use the 1-bp deletion annotations to build the deletion annotation vector. For insertions, only the impact values for the 1-bp insertion events are stored. Each entry in the annotation vector is allocated and assigned the packed annotation information that contains the corresponding coding frame and VEP annotation string from VEP annotations. Nucleotide neighborhood information is extracted from the hg38 genome sequence:

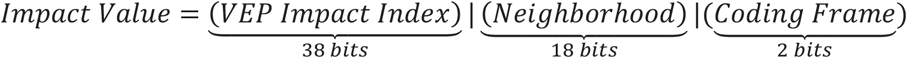

The values are concatenated and stored in a 64 bits long data type. This is a flexible representation whereby the storage can be extended with more functional information such as SIFT/Polyphen[60] scores and interspecies conservation value[60].

#### Secure SNV Variant Annotation

In the secure annotation scenario, the researcher generates the vectorized mutation array, encrypts it and sends it the encrypted data to the annotation server. The server generates the vector annotation signal as described above. The annotation of the variants is performed by multiplication of the mutation vector and the annotation vector (Fig. 2d). In the resulting vector, the entries for which there is an SNV are set to 0 and others are set to the annotation value. The server performs the multiplication over encryption and sends the results back to the user. For each position on the array that is non-zero, a variant exists. The client can use streaming operations to decrypt the downloaded data and filter out non-zero annotation entries (64-bit packed annotation information), which are unpacked according to the bit packing above. The conversion of the vector coordinates to genomic coordinates are performed while looping over the vectorized coordinates.

#### Secure Small Deletion Variant Annotation

Annotation of deletion variants is more complex than SNV annotations since deletions can have variable lengths. One way to approach this is to annotate all deletions up to a certain length, and store annotations in a vectorized annotation array for each deletion length. It is straightforward to implement this approach with the current vectorized annotation signal framework.

Rather than keeping the annotation vector for each deletion length, SVAT can also make use of the 1-base pair deletion impact signal to build the variant annotation: For deletion of nucleotides [*a, b*] (*a* < *b*), SVAT loops over all nucleotides and merges the impacts of 1-base pair deletion at nucleotide *v* in [*a, b*]. While merging, SVAT keeps track of the impact on coding sequence by counting the number of nucleotides that are deleted from CDSs. In the case that any 1-bp deletion impacts a start/stop codon, SVAT uses the 18-bit nucleotide information (coding frame) and extracts the 6-bp neighborhood to assess start/stop gain/ retainment (Fig. 2f). This is also done for the splice loss events. All retainment and loss events are annotated with the assumption that mRNAs are read from 5’ to 3’ while in translation, i.e., the reading frames are disrupted towards the 3’ end of the mature mRNA sequence after deletion and insertion events[61]. The current framework that SVAT utilizes is flexible enough to accommodate longer neighborhood sizes so regions larger than 1-codon neighborhood can be evaluated while annotating deletions.

As in secure SNV annotations, the researcher generates the vectorized 1-base pair deletion vector for all the deletion mutations that will be annotated. In this vector, any nucleotide that overlaps with deletion is set to 1 and other nucleotides are set to 0, this way the vector 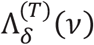 represents the deletion state of all nucleotides. These are encrypted and sent to the annotation server. The annotation server generates the vectorized annotation signal for all the 1-base pair deletions on the target regions. For this, the server annotates all the 1-base pair deletions on the target regions using VEP, then packs the impact values into the 64-bit impact array. The impact array 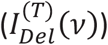 is then securely multiplied with the encrypted 1-base pair deletion vector, i.e., 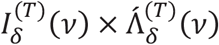. In the multiplication, any position that does not overlap with any deletion is set to 0 and other positions that overlap with a deletion contain the 64-bit packed annotation value of the 1-base pair deletion at the position. Upon receipt of the results, the researcher decrypts the annotation vector. For every consecutive position with non-zero entries, the server tracks the deletion and sets the annotation as described above.

#### Secure Small Insertion Variant Annotation

Unlike a deletion, an insertion occurs at a single location and is described by the sequence that is inserted at the location. After receiving the product of the impact vector and the encrypted insertion loci vector, 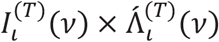, the array is decrypted. After the positions with non-zero entries are mapped back to the insertions. For this, SVAT again utilizes the 6-base pair neighborhood sequence and the coding frame at position *v* and builds the inserted junction sequence at the insertion site. This information is used to translate the codons that are inserted and the final impact on CDS, stop/start codons, and splice sites are reported.

#### Genotype Aggregation

The genotype aggregation aims to compute the frequencies of the mutations by aggregating over many samples. 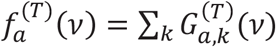, where *n*_*G*_ denotes the number of individuals in the genotype matrix, 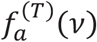 denotes the frequency of the allele *a* for the variant at vector coordinate *v* (indexed on the target regions *T*), and 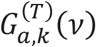 indicates the genotype of the variant for individual *k*. The entries in the genotype matrix can hold the number of alternate alleles or just the existence of mutations. Aggregation of the alternate allele counts provides the allele frequency of *a* within the sample set, as ExAC database provides. Aggregation of variant existence provides the number of samples with the mutation, i.e., similar to what genomic beacons[62, 63].

While aggregating the genotypes, it is necessary to ensure the confidentiality of the genotype matrix 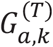 from the untrusted aggregation server, which performs the computationally heavy task of secure aggregation. To compute the above summation securely, the genotype matrix is encrypted, denoted by 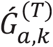 and the summation is evaluated using HE-based secure summation. Another important aspect of aggregation is to accommodate genotype matrices from multiple databases so that many samples can be aggregated together. The server needs to be able to manage multiple datasets that are encrypted with different keys. SVAT implements a proxy re-encryption protocol to convert the genotype matrices into the same key and perform the aggregation using this common key.

SVAT utilizes proxy re-encryption to securely re-code the genotype matrices (or any other type of data) so that they can be decrypted with the same secret key[64]. A trusted entity (such as NIH) is required who will perform the key management to manage the private keys necessary to generate the re-encryption keys. This is a reasonable assumption since the sensitive datasets are generally deployed and protected by entities such as NIH (e.g. database of genotypes and phenotypes -- dbGAP). When a researcher requests the aggregation service by aggregating *M* genotype matrices, the request is sent to the trusted entity, e.g. NIH., The genotypes matrices are all encrypted with different keys (as they are from different sources). The trusted entity first generates a public-private key pair and a corresponding re-encryption key for each of the matrices. The re-encryption keys are sent to the aggregation server, which re-encrypts all of the *M* genotype matrices decryptable with the same private key. Here, the aggregation server uses only the public re-encryption keys without any knowledge of the private keys. After the genotype matrices are re-encrypted, the aggregation is performed by the secure summation of the encrypted genotype matrices at every position on the target regions. The resulting frequency array, which is encrypted with the researcher’s public key, 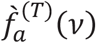, is sent to the researcher who can decrypt the frequency array and obtain frequencies.

If genotype matrices of multiple data providers are encrypted under different keys, then the server wishes to perform computation on such multi-key ciphertexts. One way to approach this is to use multi-key homomorphic encryption[65]. But the framework is still computationally intensive, making the system unsuitable for deployment in practice. Instead, we can alleviate the computational burden by the use of the conventional key switching approach of homomorphic encryption. Suppose that the key management party generates a switching key from a secret key of the input data to the common secret key and the re-encryption key is shared with the server. Then it enables the server to convert ciphertexts of genotype matrices to ciphertext decryptable with the secret key without decrypting ciphertexts.

#### SNV Aggregations

The secure aggregation of the SNVs is straightforward as they are located at one position on the vectors. In other words, the SNV frequency aggregation can be computed simply by marginalizing at every location.

#### Indel Aggregations

Unlike SNVs, the deletions cover must be tracked in each sample and aggregated. As with annotation, we make use of the 1-base pair deletions to build the aggregation of deletion of multiple nucleotides. Given a position *v*, and the 1-base pair deletion genotype matrix, 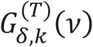; the indel of length *l*_*δ*_ is aggregated by counting the individuals *k* that satisfy the following: The deletion state of all base pairs in [*v, v* + *l*_*δ*_] are set to 1 and that the entries at (*v* − 1) and (*v* + *l*_*δ*_ + 1) are set to 0. This way, we make sure that the deletion spans exactly the coordinates in [*v, v* + *l*_*δ*_]. This procedure can effectively aggregate all the deletions that are engulfed in the target regions.

The aggregation of insertions requires explicit matching of the inserted nucleotides. This requires enumeration of all possible insertions. SVAT currently does not explicitly support aggregation of short insertion variants. However, the position at which the insertion happens can be aggregated (just by simple aggregation as for SNVs) to compute the frequency of insertion at each position.

##### Simplifications and Extensions

It should be noted that as NIH is trusted, the researcher’s keys can be generated by the NIH, too, which would minimize the computational load on the researcher, who can receive the data directly from the aggregation server. Secondly, the aggregation server does not have to store the encrypted genotype matrices. They can be discarded after the frequency vector is generated once for all the alleles and deletion lengths of interest. The above aggregation formulations can be expanded to incorporate the variant call qualities as it is done in GnomAD[49].

### Comparisons with VEP Annotations

In order to compare SVAT’s secure annotations with the plaintext VEP annotations, we simulated mutations on protein-coding genes. For this, we focused on the 10 megabase region on chromosome 1 (40mbase-50mbase) of hg38 assembly and simulated the 3 types of variants (SNVs, deletions, insertions). This region contains 6,996 exons of the 1,152 transcripts, which are used as the target regions after 100 base pair extensions. We excluded the transcripts that are tagged with “incomplete_terminal_codon_variant” and “NMD_transcript” tags. In total, the target regions covering 3,409,574 nucleotides. For all variant types, we used a 25% probability of introducing a mutation.

#### Comparison of SNV Annotations

SNVs are simulated by replacing the reference nucleotide with another nucleotide such that each non-reference allele is selected randomly with equal probability, i.e., 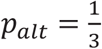. With a per-position SNV probability of 0.25, we generated 853,756 SNVs. Each mutation is annotated with VEP to generate the baseline annotations. We next annotated the simulated SNVs using SVAT. We finally compared the annotations in terms of matches between assigned impact strings for each variant. As expected, VEP and SVAT have yielded exactly the same annotations.

#### Comparison of Deletions

We described 2 approaches that deletions can be annotated. The first approach is using deletion-length specific annotation vectors (and corresponding mutation vectors). Similar to SNV annotations, this approach will enable us to exactly replicate the VEP annotations as it explicitly encodes the impact strings into the annotation vector for deletions of every length.

The second approach uses the 1-bp deletion annotation vector and 1-bp deletion variant loci vectors. As described earlier, the impact of each deletion whose length is greater than 1-bp needs to be translated using the 1-bp deletion annotations. The impact values assigned by SVAT may, therefore, not perfectly match VEP annotations. To evaluate the mismatches systematically, we simulated 853,145 deletion variants whose lengths are randomly selected (uniformly distributed with [1,10]). The simulated deletions are annotated using VEP. Then we simulated the secure annotation by SVAT where the 1-bp deletion annotation signal is multiplied with 1-bp variant loci vector and the product vector (i.e., annotated vector) is translated to generate the annotation for all the deletions.

While comparing the annotations, we first focused on the 6 impact values that are assigned by VEP whose impact strings are classified with the HIGH impact category. These are (1) frameshift_variant, (2) splice_acceptor_variant, (3) splice_donor_variant, (4) stop_gained, (5) stop_lost, (6) start_lost. Next, we compared the high-impact annotations that are assigned to the variants by VEP and by SVAT. For each high-impact annotation assigned by SVAT (VEP), we counted the frequency of mismatching annotations assigned by VEP (SVAT).

#### Mismatch of HIGH Impact Category

We first counted the number of mismatches in the HIGH impact category where we found the number of annotations where SVAT and VEP did not assigned a high impact annotation. We found that out of 2.7 million matching annotations (Per variant and transcript), 2,068 (Less than 0.1%) annotations contain a mismatch where SVAT or VEP assigned one of the HIGH impact annotations while the other did not. We next analyzed the mismatches for each HIGH impact category. Figure 3 shows the comparison of high-impacting terms. We describe the mismatching annotations and justify the annotations assigned by SVAT. We believe this comparison is reasonable since annotation of indels is not definitive and different methods annotate indels differently[66].

**Figure 3.**
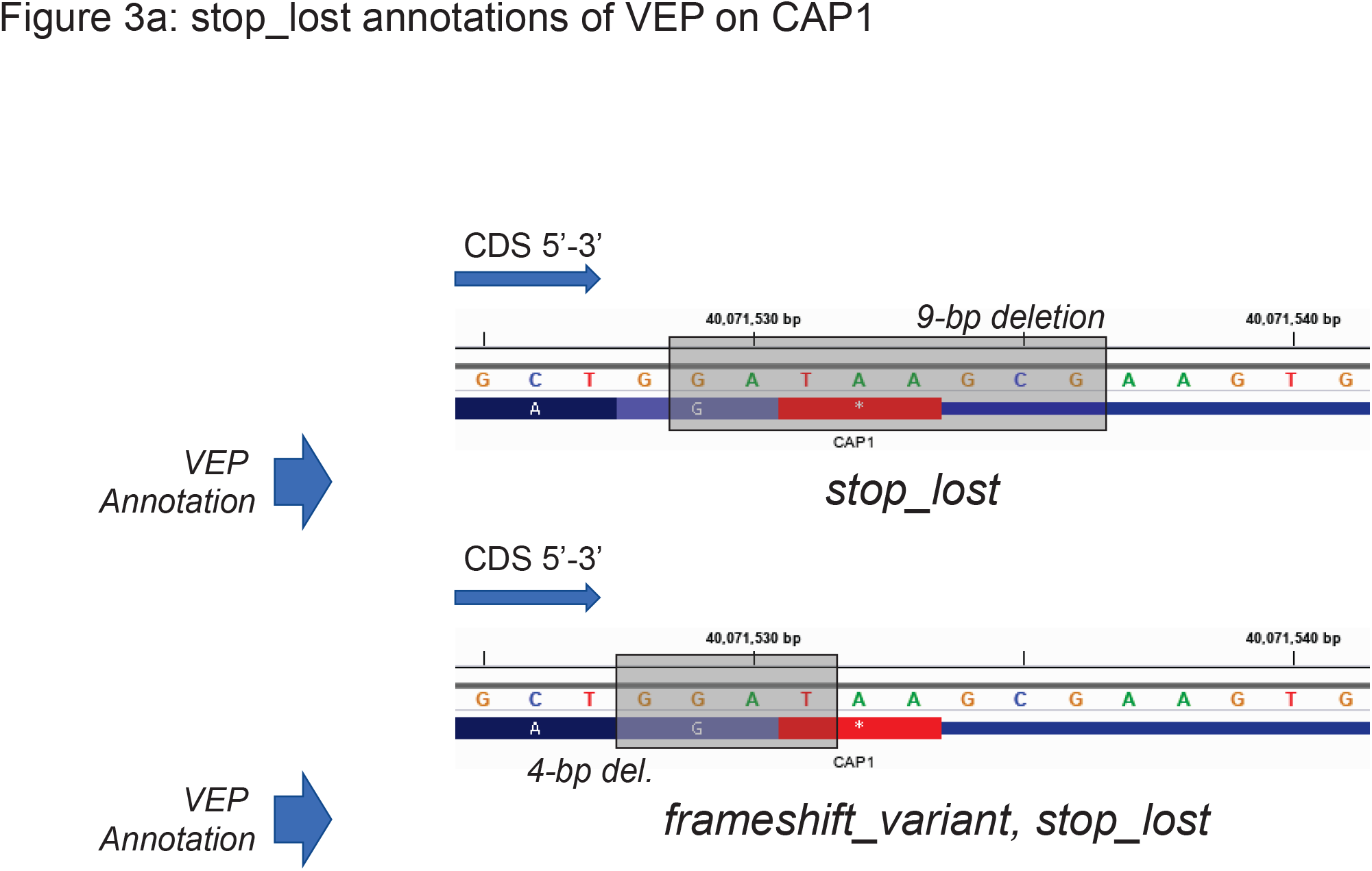
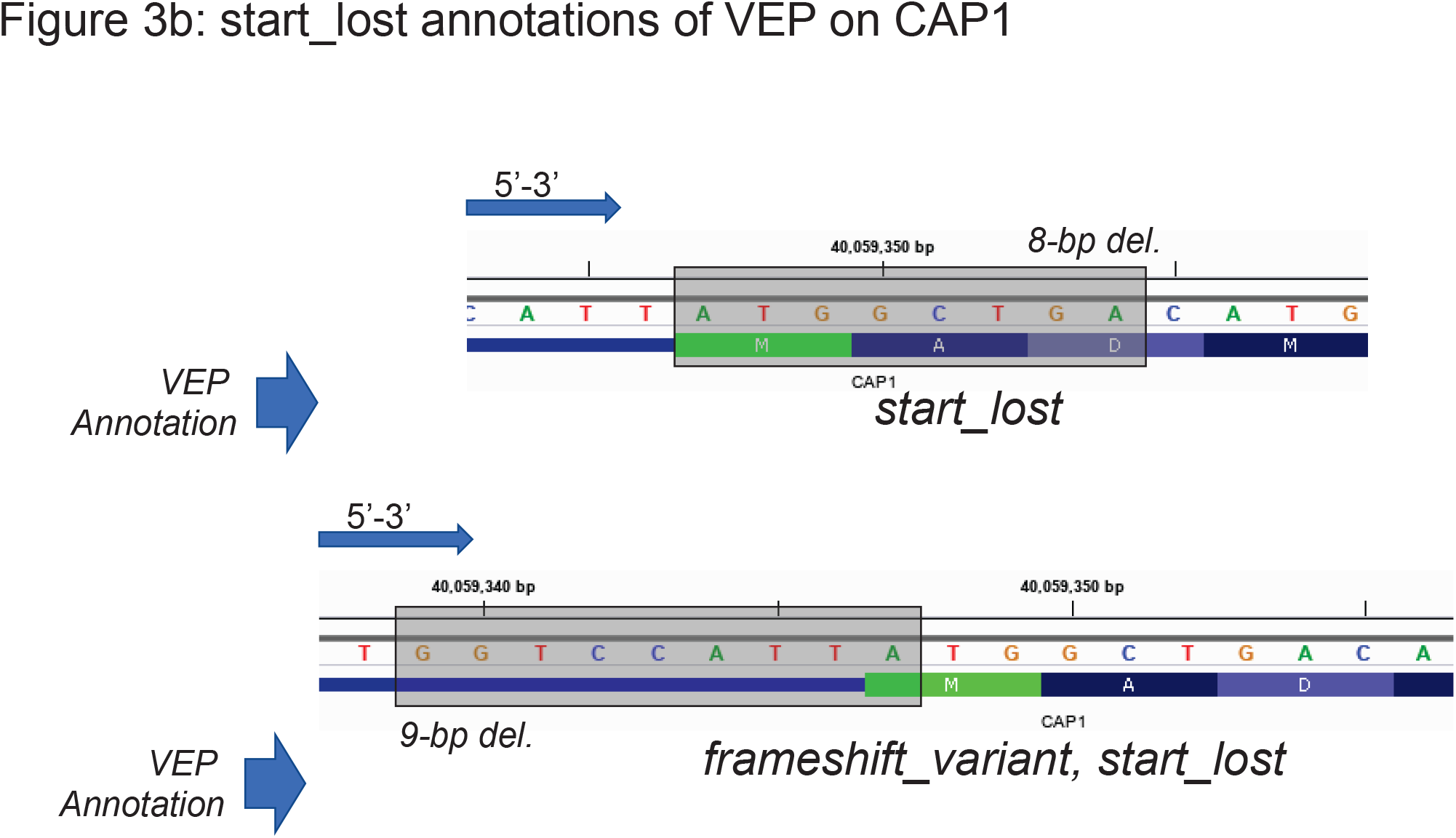
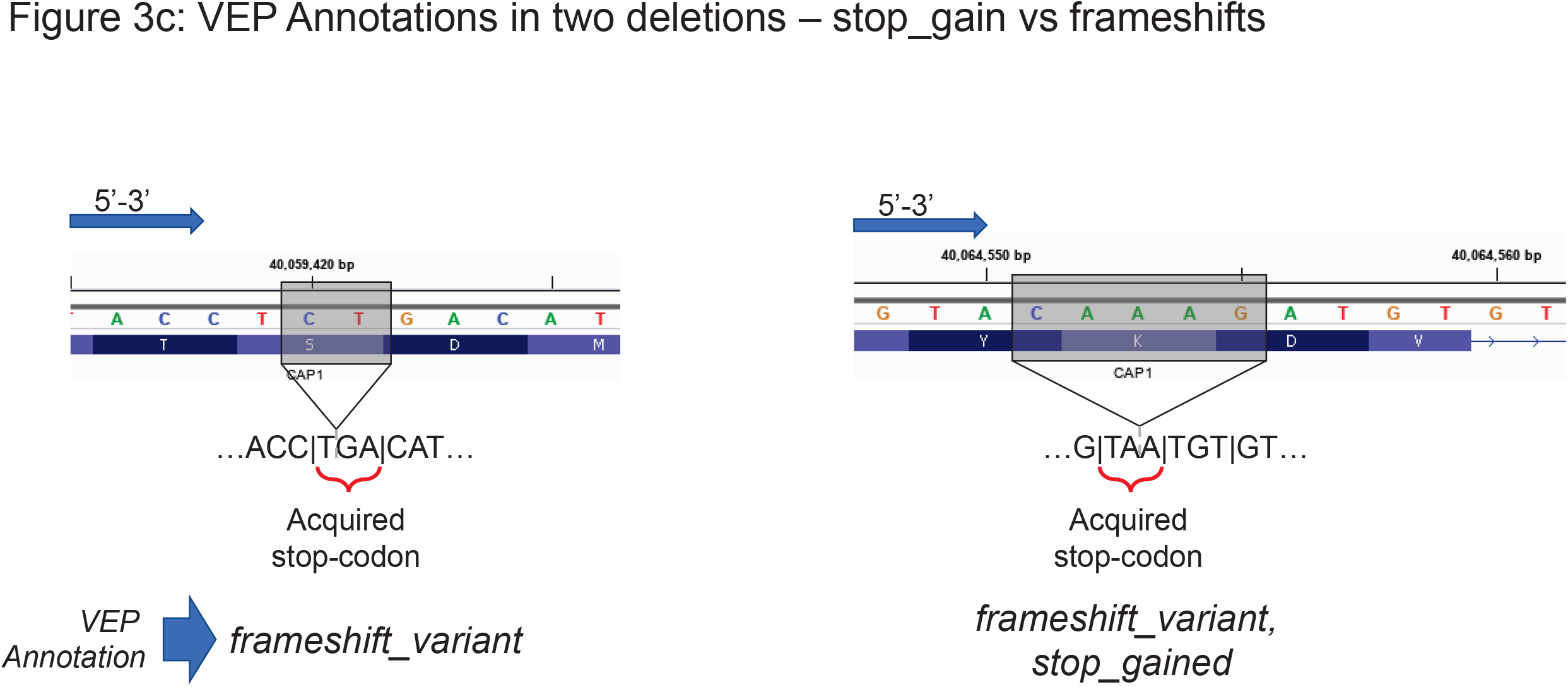
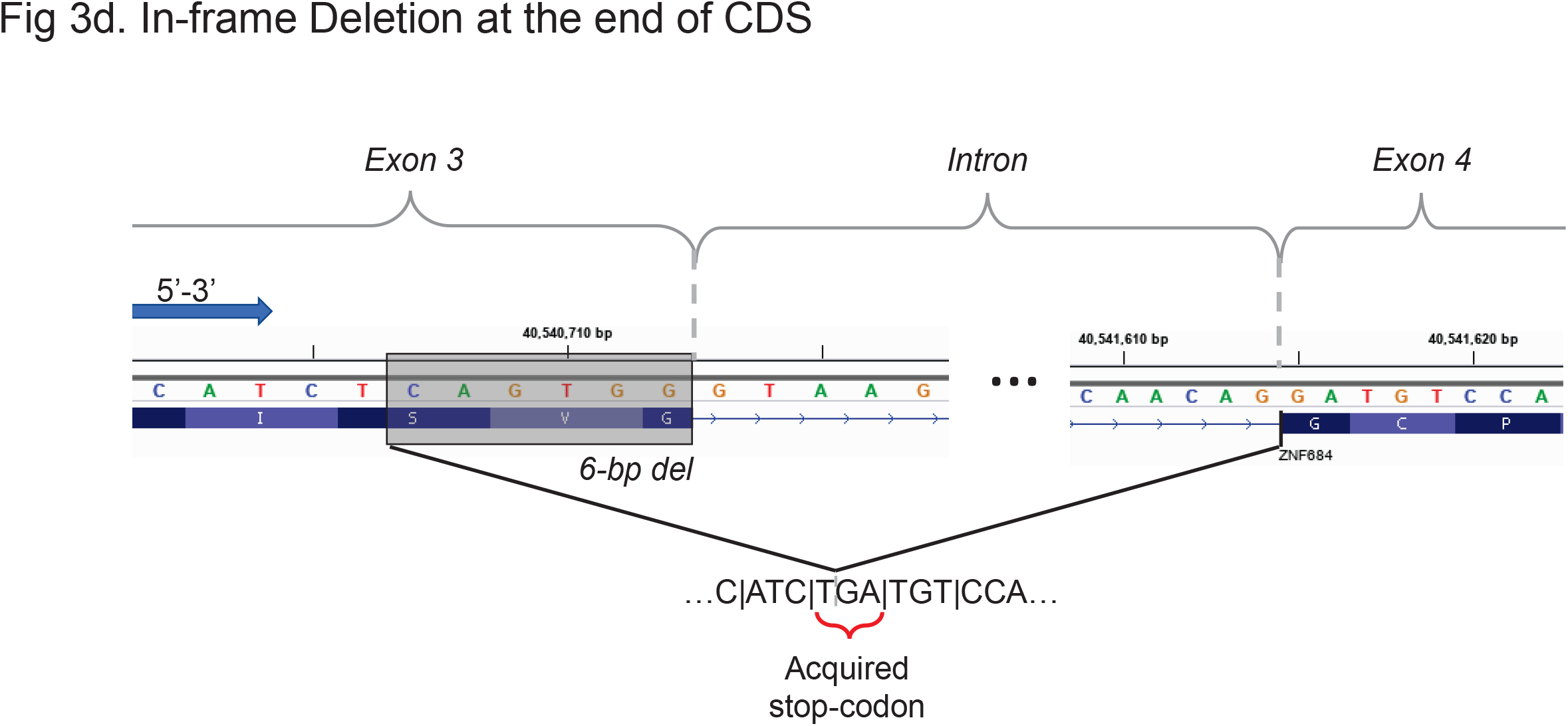
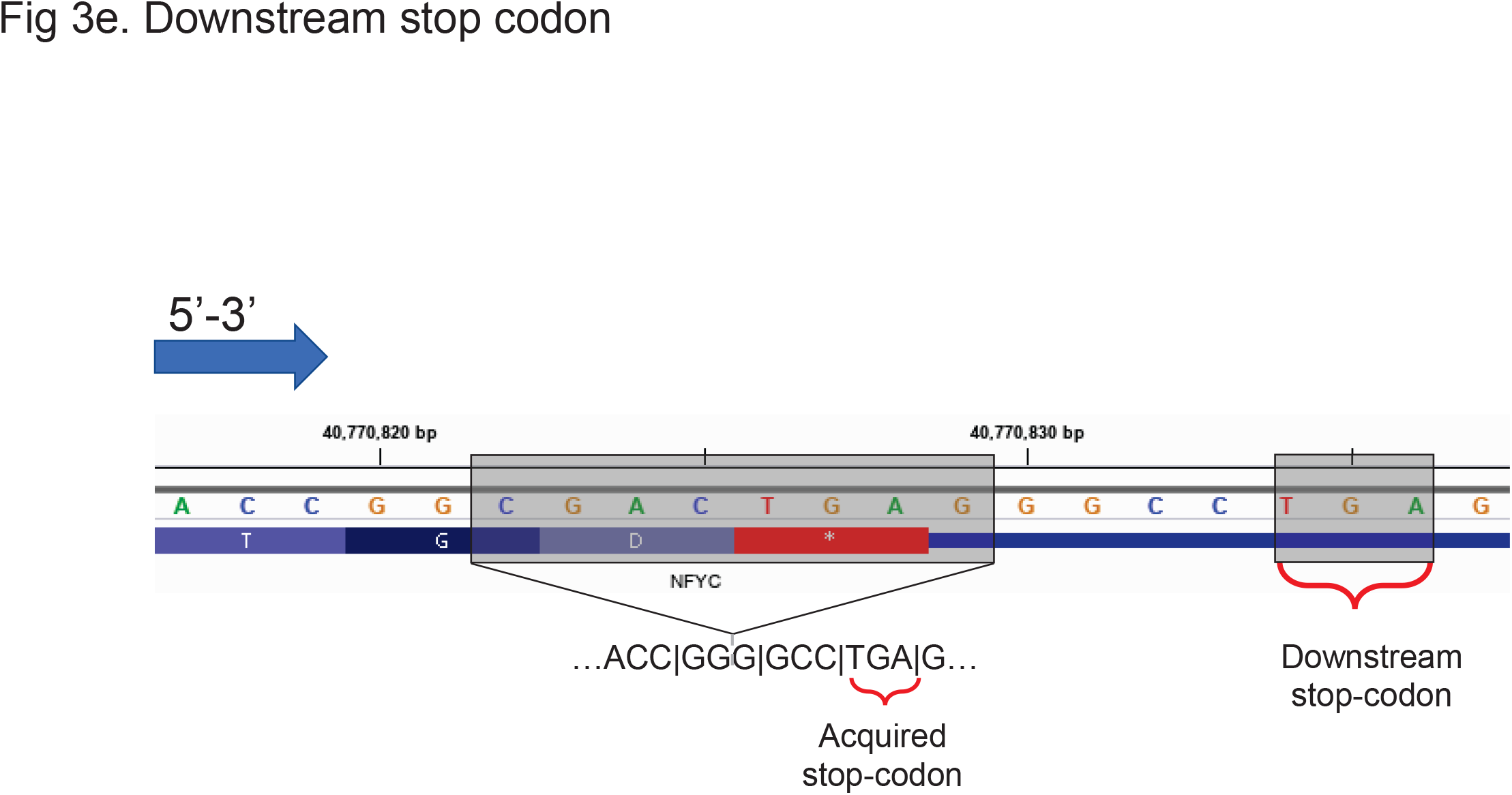
Examples of mismatches between SVAT and VEP while annotating deletions. (a) Two examples of deletions that impact CAP1 gene as they are annotated by VEP. 9-bp deletion that engulfs the stop-codon (shown in red) are annotated by VEP. This variant is annotated by VEP only as “stop_lost”. On the other hand, the 4-bp deletion that deletes only 1 base pair from the stop-codon is annotated as “frame_shift, stop_lost”. (b) Similar to (a), the start-codon impacting variants are annotated by VEP as “start_lost” for an 8-bp deletion (top) and as “start_lost, frameshift” for a 9-bp deletion (bottom). VEP does not assign “frame_shift” term when the start-codon is engulfed in the variant. In both of these cases, SVAT assigns both frame_shift and start_lost terms. (c) Examples of two deletions that are discordantly annotated by SVAT and VEP. For both deletions, SVAT annotates by “frameshift_variant, stop_gain” whereas VEP annotates the 2-bp deletion (left) as frameshift and the 5-bp deletion (right) as “frameshift_variant, stop_gain”. 2-bp deletion (left) also causes a stop-gain at the junction where frame contains a “TGA” stop-codon. This is detected by SVAT while VEP does not report the stop-gain. It should, however, be noted that these deletions are both in HIGH impact class and would be picked up as candidates in downstream analyses. (d) An example of a stop_gain that is detected by VEP and missed by SVAT. The deletion that ends right at the splice junction site creates a junction with a stop codon. Both mutations are in HIGH impact class as this is also a frameshift mutation. (e) Another example of a downstream stop-codon that is missed by SVAT and picked up by VEP. The 8-bp deletion engulfs stop-codon and creates a new frame such that a downstream stop-codon (“TGA” stop codons as shown in figure) is created within the new coding frame. This mutation is marked as HIGH-impact by VEP and by SVAT.

#### SVAT Specific frameshift annotations (Missing in VEP annotations)

The most frequent mismatch in annotations are variants that are annotated as splice_acceptor or splice_donor by VEP whereas SVAT assigns them as frameshift mutation, in addition to splice_acceptor or splice_donor. Upon inspection, we found that VEP’s annotations are disjoint for these classes, i.e., splice_acceptor and splice_donor annotations do not overlap with frameshift annotations.

The stop and start loss are the two other mismatching VEP annotations where SVAT additionally assigns frameshift annotations. These mutations are also annotated with stop and start loss impacts by SVAT, in addition to frameshift. We also observed that VEP annotates start and stop losses as frameshift only when the deletion is exclusively in the coding sequence and impact a start or a stop codon by 1 or 2 nucleotides (Fig 3a). Thus, SVAT performs a more complete annotation of these mutation classes.

Similarly, numerous mutations SVAT annotates with frameshift and stop_retained are annotated only as stop_retained by VEP. We inspected several of these mutations and found that Annovar annotates these mutations as frameshift as well. A small fraction of these mutations are found to be downstream stop codons that VEP traces. For these mutations, SVAT does not assign stop_retained impact. This is because SVAT analyzes only one codon that the deletion impacts. This is one of the current limitations of the default parameters of SVAT that uses 3 base pair neighborhood sequence and can be mediated by increasing the neighborhood size.

We also would like to highlight the cases of start_retained and start_lost impact classes. VEP annotates some of start_lost mutations with start_retained. We chose to exclude the start_retained variants exclusively because the start codon is surrounded in the upstream by conserved binding motifs that are used for translation initiation, such as the Kozak motifs[67]. This does not apply to stop_retained class since stop codons do not require surrounding sequence motifs to terminate translation, unlike start codons (Fig. 3b).

#### SVAT specific stop-gain annotations

We next focused on SVAT specific stop-gain annotations. Among these, most of them are annotated as frameshift by VEP. All of these mutations are also annotated as frameshift with SVAT. While we inspected numerous cases, we could not identify a specific reason why VEP chooses not to annotate these variants as stop-gain (Fig 3c). We also found that VEP does not assign stop-gain to some of the inframe deletion mutations that seem to introduce early stop-gains in the coding sequence. Previous studies have pointed out the discrepancies among annotation tools about assignment of stop-gain mutations[66].

#### SVAT specific stop-lost annotations

We observed that majority of the mutations that are annotated as stop-lost only by SVAT are annotated as stop-retained by VEP. These are annotated as stop-retained by SVAT as well (in addition to stop-lost). Upon inspection, we found that the main reason for this discrepancy is that VEP assigns stop-lost and stop-retained annotations disjointly.

#### VEP specific stop-gain annotations

There are mutations that manifest at the splice sites such that the deletions at the end of CDS create a stop-codon on the processed mRNA sequence after splicing. This case is currently not handled by SVAT yet, and represents another limitation (in addition to the tracking of the downstream stop-gains) of SVAT (Fig. 3d,e). This case can be handled by keeping track of the neighborhood coding sequence, in addition to the genomic neighborhood.

#### Comparison of Insertions

For comparison of insertions, we performed a simulation with a 25% chance of having an insertion at any position. For any insertion, the length is selected uniformly between 1 and 10 base pairs and inserted sequence is generated as a random string of {A,C,G,T} nucleotides. We generated 854,639 variants on the target and annotated them with SVAT and VEP. It should be noted that the translation phase of insertions only requires translating the inserted sequence and computing the frame. Out of the 2,352,611 annotations that are matching in variant and transcript among VEP and SVAT, we identified only 246 annotations exhibit a mismatch at the high impact category, i.e., either one of the methods provide a HIGH impact annotation. When we analyzed each impact string (as for deletions), we observed a similar pattern where SVAT-specific stop-gain and stop-loss annotations are matched by frameshift and stop-retained annotations that are assigned by VEP, as with deletion cases. The stop-gain that manifest by the insertions at the ends of exons is again missed by SVAT (as expected) (Fig 3d,e).

Overall, these results indicate that SVAT has comparable and more exclusive annotations compared to VEP. We do not provide these results as SVAT performing more accurately than VEP because SVAT is based on VEP annotations. In addition, the default parameters of SVAT may exhibit several limitations for a small number of events, namely (1) downstream stop-codons and (2) indels at the end of CDSs.

### Secure Annotation Resource Requirements

We next analyzed the resource usage of SVAT’s secure annotation module. We divide the annotation task into 5 consecutive steps: (1) variant loci vectorization, (2) Encryption of vectorized vector, (3) Multiplication with vectorized annotations, (4) Decryption, (5) Translation. We report the time, memory and disk space usage for each step. Table 2 shows the requirements for annotation of 3,409,574 vectorized positions (chromosome 1:40,000,000:50,000,000) and simulated 50,000 deletion variants.

**Table 2:** The time, memory, and disk space requirements for SVAT variant annotation for 50,000 deletion variants on target regions covering 3.4 megabases.

|  | Time (Seconds) | Memory (Megabytes) | Disk Space (Megabytes) |
| --- | --- | --- | --- |
| (1) Variant Loci Vectorization | 1.51 | 20.02 | 1.3 |
| (2) Encryption | 0.054+ 0.030=0.084 | 43.90 | 43.90 |
| (3) Secure Multiplication | 0.040298 | 429.83 | 43.90 |
| (4) Decryption | 0.100407 | 429.83 | - |
| (5) Translation | 7.79 | 165.49 | 4.2 |
| Total Client/Total Server | 10.24 /0.040298 | 165/429 | 4.2/43.9 |

SVAT requires less than 10 seconds of computation time. The timing requirements on the annotation server are around 1 second in our benchmark. The maximum main memory usage among the 5 steps is less than 0.5 gigabytes. These requirements are fairly modest compared to the general pipelines. In general, however, the step of (1) vectorization, (2) encryption, (4) decryption, and (5) translation will be performed at the client, who may have restricted computational resources. Among these steps, translation of the annotations requires the highest but fairly modest time and memory, which can be feasibly implemented into commodity computers. The disk space usage is important as disk space may be limited at the client. We observed that the encrypted data for 3.4 million positions could be stored in approximately 43 megabytes of encrypted data, i.e., the client needs to submit 13 bytes/vector position for the encrypted data, which is related to the expansion rate of encryption and the data representation. This requirement will be reflected in the disk space requirement and the network usage. As the encrypted data size and usage increases linearly with the number of vectorized positions, we can estimate the resource requirements for secure annotation of protein-coding genes. The gene-specific target regions cover 48 megabases. From Table 2, we can estimate that the maximum memory requirement (for gene-specific target regions) is less than 8 gigabytes for the server and 3 gigabytes for the client. In terms of timing, the whole gene-specific target regions can be annotated in less than 3 minutes (180 seconds). We do not report the time for key-generation steps as they are substantially smaller than the reported times.

### Resource Requirements of Genotype Aggregation and Re-Encryption

Genotype aggregation for SNVs is performed by marginalization (i.e. sum) over the rows of the genotype matrix. To evaluate the time/memory and disk space requirements, we focused on the aggregation of the alternative allele frequencies over 1000 individuals using the genotype data from the phase3 of 1000 Genomes Project[2]. As the target regions, we focused on the exons of 1000 genes on chromosome 1 that covered 6.6 megabases of nucleotides. We divided the aggregation task into 3 different steps and report resource requirements for each step: (1) Encryption, (2) Aggregation, (3) Decryption. For the aggregation task, SVAT aggregates the genotypes allele-by-allele (i.e., the genotype matrices corresponding to allele A, C, G, T), which can help decrease memory usage. Table 3 shows the resource requirements for genotype aggregation for each allele. The time and memory requirements are measured using 24 threads. The longest time for the aggregation task is spent in encryption: 96 seconds are spent for encrypting the 6.6 million-by-1000 genotype matrix for one allele. The disk space usage for one allele is around 83 gigabytes (approximately 13 bytes/position/sample). Aggregation takes around 2.56 seconds and uses 107 gigabytes of memory. Finally, the decryption takes around 0.2 seconds of time and decrypts the 6.6 million long aggregated counts of the alleles at each position using 0.34 gigabytes of memory.

**Table 3:** The resource requirements for aggregation of 6.6 megabases-by-1000 individuals genotype matrix for one allele.

|  | Time (Seconds) | Memory (GiB) | Disk Space (GiB) |
| --- | --- | --- | --- |
| Encryption (Client) | 96.26 | 17.94 | 83.01 |
| Secure Aggregation | 2.56 | 106.98 | - |
| Decryption (Client) | 0.17 | - | 0.34 |

Our results indicate that the aggregation task can be more computationally intensive compared to the annotation task. This is reasonable since the genotypic data used in the aggregation task is much larger than the variant locus information that is used for annotation tasks. Our results also provide evidence that the vectorized representation can enable large optimizations and increased speed in the secure annotation and aggregation tasks; these are enabled by the streaming operations that the new HE schemes take advantage of such as data packing and vectorization operations[33]. There is, however, an accompanying bottleneck on the disk space and memory usage of the vectorized representations.

We finally evaluated the resource requirements of the re-encryption-based pooled aggregation of genotype matrices, which enables aggregating genotype matrices that are obtained from different sources (different private keys, Fig. 1). For this, we tested the pooling of two genotype matrices where the first matrix (*M*_1_), contains the genotypes at 6.6 million vectorized positions for 500 samples and second matrix (*M*_2_) contains the genotypes for 1000 samples, i.e., pooling of 1500 samples from two databases. We evaluated the 4 steps: (1) Encryption of each matrix in their own keys, (2) Re-encryption into the same private key and pooling of the matrices, (3) Aggregation of the pooled matrix, (4) Decryption of the results. Table 4 shows the time requirements of these operations for the pooling of the two datasets using 24 threads.

**Table 4:**
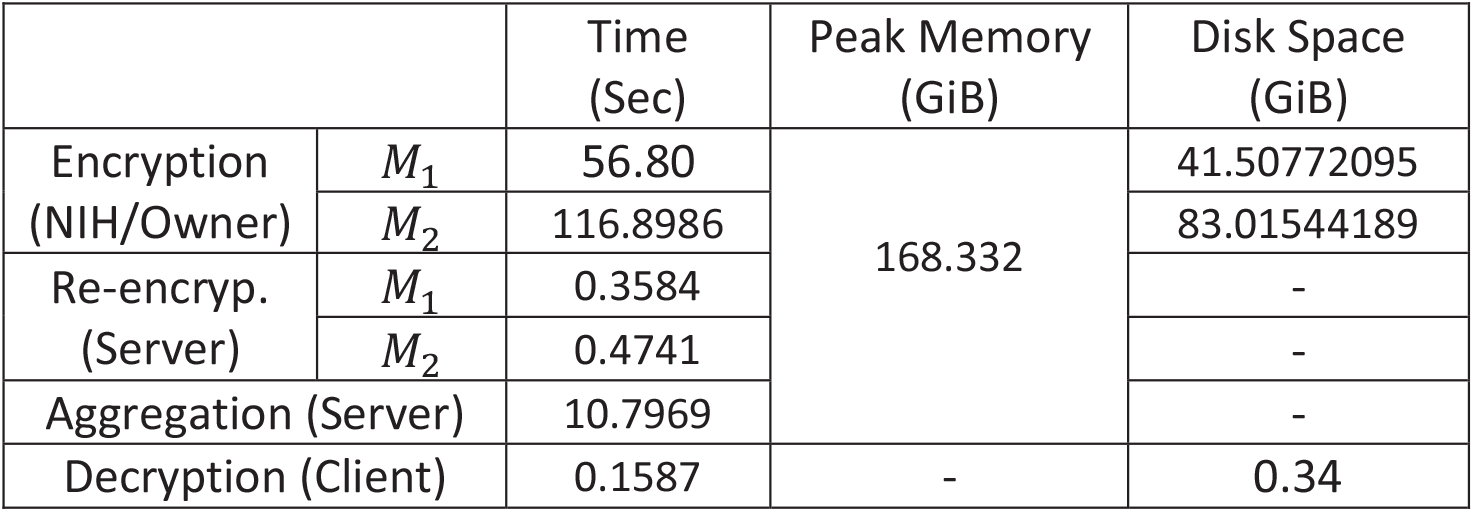
The resource requirements of aggregation tasks. Encryption is performed at the owner or the trusted entity, i.e., (NIH). The encrypted data is sent to the aggregation server. After the client requests the aggregation, the trusted entity generates the re-encryption key and sends it to the server. The genotype matrices are re-encrypted and aggregated by the server.

|  |  | Time (Sec) | Peak Memory (GiB) | Disk Space (GiB) |
| --- | --- | --- | --- | --- |
| Encryption (NIH/Owner) | $M_1$ | 56.80 | 168.332 | 41.50772095 |
| | $M_2$ | 116.8986 | | 83.01544189 |
| Re-encryp. (Server) | $M_1$ | 0.3584 | | - |
| | $M_2$ | 0.4741 | | - |
| Aggregation (Server) |  | 10.7969 |  | - |
| Decryption (Client) |  | 0.1587 | - | 0.34 |

The most time-consuming step is encryption, which took slightly more than 116 seconds for *M*_2_. The re-encryption operation takes less than 1 second for both matrices. This is reasonable since the encryption can be performed once at the trusted entity of the owner and re-encryption can be performed many times on the aggregation server. The aggregation operation takes less than 10 seconds and finally, the decryption operation takes less than 1 second. Disk space usage is moderate with 83 gigabytes of data that needs to be stored for the full genotype matrix. It is, however, not necessary to store the genotype data for aggregation operations. The server can generate the encrypted aggregations for each matrix, then store the encrypted vector containing the aggregated allele counts, which takes much smaller space than the genotype matrices.

## Discussion

We presented SVAT, a method for secure annotation of variants and secure aggregation of genotypes from multiple databases. The increasing number of genomes frequently makes it necessary to perform batch annotation and aggregation of variants by outsourcing these operations with programmatic or through user interfaces on web servers. SVAT makes use of homomorphic encryption to provide confidentiality to the genotype data and variant loci. The proposed framework makes use of a novel vectorized representation of the variant loci to protect the variant loci information. This new representation is utilized commonly for annotation and aggregation tasks. This can enable building more complex variant analysis pipelines with allele frequency and variant impact filters. An alternative method that is similar in flavor is “private-set-intersection” (PSI) [68, 69], where two entities can securely intersect of their confidential datasets without revealing any information other than only the intersecting elements. In particular, our approach is similar to the PSI implemented in Chen et al.[70] albeit not in the genomic data analysis context. Chen et al. the authors design a multiplication statistic that is computed for each data point in the query dataset (i.e., researcher’s variant dataset) to compare every data point in the server’s dataset using an HE-based secure subtraction and multiplication operations. This encrypted data is returned to the client who decrypts and identifies the 0 entries in the returned vector, which points to intersecting elements. As Chen et al. describe, this approach works with small query datasets. Our approach limits the search space by using target regions and similarly enumerates all possible mutations and vectorizes the operations. A simple PSI query must be combined with the extra information so that annotation information is returned, which, we believe, is not a simple task. On the other hand, we assume that the server’s annotation dataset is not confidential, which is a strict condition that is satisfied with a PSI search. In addition, the statistic used by Chen et al. can be modified to accommodate plaintext server data.

There are alternative approaches that can enable the confidentiality of the genetic data. For example, just for the annotation task, the client can download the vectorized annotation data and perform annotation locally without ever having to upload the data. This would be similar to the “share models, keep data” approach[71], where the researchers can download the annotation vector and extract annotation information for their variants without ever sending the variants from their studies. This is, however, a different paradigm than what is undertaken in this study. SVAT aims to enable a data sharing paradigm where the encrypted data stays with the untrusted entity and can be re-used by other HE-enabled pipelines. The resharing can be enabled by the proxy-re-encryption module of SVAT. We acknowledge that there is more work to be done to render the framework more practical so that it can be run on whole genome scale. While the computational framework can accommodate any annotation and aggregation task, there are several limitations of SVAT that are used to bound the computational and memory cost. First, the target regions are used to decrease the vectorized data size. In order to focus on the most impactful mutations, SVAT uses the protein-coding exons and surrounding regions as the targets. Second, SVAT can decrease the time and memory usage for the annotation of deletions by making use of an annotation vector that contains the 1-bp deletions and making use of this to translate the impact of deletions that span multiple nucleotides. This approach is exact for the highest impact that are assigned to each deletion.

## Conclusions

Variant annotation and genotype aggregation are two integral components of genomic data analysis pipelines. For instance, the software pipelines that perform variant association tests use variant annotations to classify variants based on their impact. Similarly, genotype aggregation is used for classifying and filtering genetic variants with respect to population-level allele frequencies. This is an integral step to estimating the selection pressure and detecting potentially disease-causing variants. SVAT can be integrated into these analysis pipelines to securely store and analyze datasets. The presented framework can be used in other context and potentially has overarching impact on other prospective privacy-aware method development efforts.

## Methods

We present the details of the annotation and aggregation workflows.

### Variant Annotation

Target regions are provided as a BED file with name and strand entries. The default target regions comprising the extended CDS coordinates are included with SVAT for hg38 genome and Gencode v31 annotations[54]. The target regions can be modified using a GFF/GTF formatted gene annotation file with exon, CDS, and UTR entries present. The users can also specify the number of extension of each element, i.e. *l*_*ext*_, to include the functional elements around exons/CDSs using “transcript-specific” and “gene-specific” target regions as described in the previous Section. The input variant coordinates to be annotated are taken as a VCF file by default. SVAT processes VCF files to generate the vectorized variant loci signals.

### Vectorized Representation of the Variant Loci

The vectorization enables protection of variants that fall on the target regions. The exon start/end coordinates are extracted for the protein coding transcripts. Each CDS is extended by *l*_*ext*_ to include variants that may impact splicing. The target regions are described with pairs of coordinates:

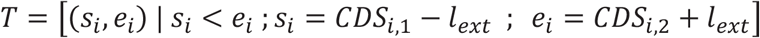

where *T* denotes the sorted list of target regions, *s*_*i*_ and *e*_*i*_ denote the genomic coordinates for the start and end, respectively, of the *i*^*th*^ target region, and *CDS*_*i*,1_ and *CDS*_*i*,2_ denote the start and end coordinates of the *i*^*th*^ CDS. Each target region corresponds to a CDS and is assigned to an element, whose id is known, e.g. transcript/gene name. *T* is a sorted list and the sorting cannot be changed in the course of annotation for the mapping from genomic to vector coordinates is used multiple times. While the actual sorting of the regions does not have any specific importance, position-based sorting will make conversion of genomic coordinates to target regions efficient using optimized searching algorithms. By default, SVAT first sorts the target regions first by start position, then sorts with respect to the element identifier, i.e., transcript/gene identifier. To denote the *i*^*th*^ target region, we use *T*_*i*_, i.e., *T*_*i*_ (*s*_*i*_, *e*_*i*_). Also, we denote the total nucleotides covered by target regions by *l*_*T*_:

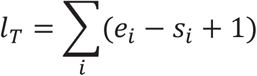

Next, the sorted target regions list is used to build the vectorized signals. For this, SVAT allocates an array of length that is equal to the coverage of the target regions *T*:

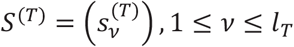

where *S*^(*T*)^ denotes the vectorized signal, whose coordinates correspond to the target regions *T*. Every position on *S*^(*T*)^ holds a value that corresponds to a position on the target regions. The mapping between the position on the vectorized signal and the sorted target regions list is done by tracking the target regions. It is necessary to identify the target region index, and then a position index on the target region so that we define exactly where the vector position maps to on the genomic coordinates. First, we identify the leftmost sorted target region whose cumulative coverage is smaller than *v*:

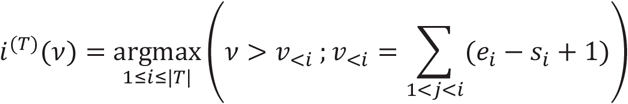

where *v*_*<i*_ indicates the total cumulative number of nucleotides that are covered by the target regions *T*_1_, *T*_2_,…, *T*_*i*_, and *i*^(*T*)^(*v*) indicates the target region index (*i*^(*T*)^(*v*) < |*T*|) that the vectorized position *v* corresponds to. The above formula, although complex looking, simply states that we start from the leftmost target region and track the total number of nucleotides that are covered by the sorted target regions list. The largest region’s index such that *v*_*<k*_ *< v < v*_*<*(*k*+1)_ where *k i*^(*T*)^(*v*). This can be performed by a binary search over the sorted target regions list. After the target region index is identified, we identify the position that corresponds to the target region as:

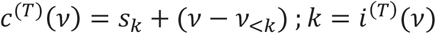

where *c*^(*T*)^(*v*) indicates the genomic coordinate of the vectorized position *v* when it is mapped onto the target regions *T*. The mapping of the target coordinates onto the vector coordinates can be performed similarly: *v*^(*T*)^(*i, c*) *= v*_*<i*_ + (*c* − *s*_*i*_). It should be noted that we can only map the coordinates of the targets to vectors in a 1-1 fashion. The genomic coordinates can be mapped to multiple target regions (and to multiple corresponding vectorized positions). To identify all vector coordinates that map to a genomic coordinate (*pos*), we identify all the target regions that overlap with this position:

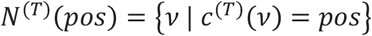

where *N*^(*T*)^(*pos*) denotes set of vector coordinates that map to genomic coordinate *pos*. To implement the above search, target regions whose genomic coordinates overlap with *pos* can be identified then formula for *v*^(*T*)^(*i, c*) can be used for each overlapping target region.

### Vectorized Mutation Loci

An array of length *l*_*T*_ that is indexed by the vectorized coordinates (with respect to a sorted target region list) is used to store the variant locus information. Given a variant allele *a* (*a* ∈{*A, C, G, T, δ, L*} (the alternate alleles of an SNV, 1-bp deletion, and 1-bp insertion), variant loci are mapped onto the vectorized coordinates as described previously. Next, for each allele, an array of *l*_*T*_ is allocated and each array entry whose vectorized position overlaps with a variant is set to 1. All other positions are set to 0 (Fig 2d):

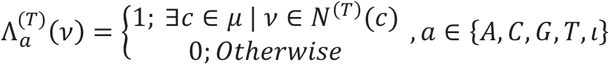

where 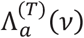 denotes the variant loci array for allele *a, µ* denotes the set the genomic coordinates of the mutations. The above equation simply describes that the position *v* on the mutation loci array is set to 1 if there exists a mutation with allele *a* whose genomic coordinate maps to *v*, i.e. *v* ∈ *N*^(*T*)^(*c*). For the deletions, every position on the deletion is set to 1:

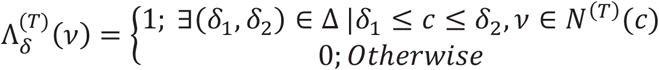

where Δ denotes the list of start-end coordinates for the deletions and *δ*_1_, *δ*_2_ denote the coordinates of the first and last nucleotides that are deleted in the deletion. To set the array, we can simply iterate over all deletions in Δ, then for every genomic position *δ*_1_≤ *c* ≤ *δ*_2_, find the mapping vector coordinates and set the vectorized deletion loci signal to 1.

### Encryption of the Vectorized Variant Loci

For simplicity, we assume that *l*_*T*_ is divisible by the plaintext length bound *l*. Then an array of length *l*_*T*_ is divided into plaintext vectors of size *l*, each of which is encrypted using the public key of the cryptosystem. Otherwise, an input array is divided in a way that as many sub-vectors as possible have the maximal size *l* and the only last vector is chosen to have a smaller size than others. To be specific, the encryption procedure results in *r* ⌈*l*_*T*_*/l*⌉ ciphertexts.

### Vectorized Annotation Information

SVAT uses an array of length *l*_*T*_ to hold the annotations of all mutations on the target regions. This is very similar to the mutation loci vector, except that for each position *v*, the array stores a 64-bit entry that packs the annotation information.

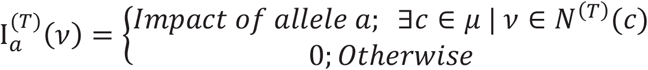

The impact value indicates the impact of the mutation located at vector position *v* with allele *a*, and *µ* denotes the set of genomic coordinates (i.e., indices) for all mutations. The impact information is retrieved from Variant Effect Predictor, VEP, by default. SVAT packs the impact information and the nucleotide information around the mutation locus. The packed impact information contains:

1. Coding frame (2 bits): This is the coding frame of a mutation that is a value in {0,1,2}: The frame is converted to a 2-bit value and used as is.
2. 6-base pair nucleotide neighborhood (18 bits): SVAT assigns 3-bits values (A: 000, C: 001, G: 010, T:011, N:100) to each nucleotide and codes the surrounding nucleotide sequence into 18 bits. The sequence of bits is concatenated and used as the neighborhood sequence information.
3. The assigned impact string identifier assigned by VEP (38 bits): The bitmap that describes the impact values of the mutation out of the 38 different impact values that VEP assigns to each mutation. The bitmap is generated by setting the impact values to 1 for every impact string assigned by VEP. For every annotation, the index of the impact string in the bitmap is set to 1.

The annotation server generates the vectorized annotation signal. In order to generate the vectorized annotation information, VEP is run to generate the annotation of all the nucleotides (and all alleles) of the mutations on the target regions. This is performed for 4 alleles of SNVs, 1-bp deletions, and 1-bp insertions on all positions on the target regions. The output of VEP can be directly piped into SVAT to convert the VEP annotations into the vectorized annotation signal. SVAT first maps the position to the vectorized coordinates. Next, the coding frame information and the 38-bit impact bitmap are updated using the assigned frame and the list of impact strings. SVAT finally extracts the 6-bp neighboring nucleotides from the genome sequence (by default, hg38 genome sequence) and builds the 58-bit annotation vector value. The annotation value is assigned to all of the vector coordinates that map to the mutation’s genomic coordinates. The packed impact value can be formulated as:

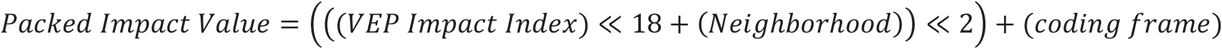

The whole operation for building the vectorized annotation signal is a compute-intensive task. However, this needs to be performed at the annotation server. We assume that the annotation server has large computational resources and can perform this operation. In addition, the annotations need to be vectorized once for every gene annotation.

### Secure SNV Variant Annotation

After the variant loci and the annotations are vectorized, the annotated variants vector is computed as the multiplication of the variant loci vector and the annotation vector (Fig. 2d). The results from this product is a vector of length *l*_*T*_:

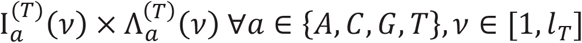

In this vector, the entries for which there is no variant are set to 0 (since 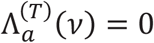 when no mutation maps to *v*) and others are set to the annotation value at the position.

Suppose that for *j* = 1,2, …, ⌈*l*_*k*_*/l*⌉, a plaintext vector 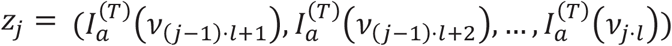 is encrypted into a single ciphertext *ct*_*j*_. The server performs the constant-ciphertext multiplication with the ciphertext of the annotation vector and the corresponding variant loci vector as follows:

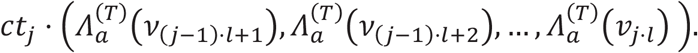

Then the results are sent back to the user. For each position on the array that is non-zero, a variant exists. Streaming operations are used by the client to decrypt the downloaded data and filter out non-zero annotation entries (64-bit packed annotation information), which are unpacked according to the bit packing above. The conversion of the vector coordinates to genomic coordinates (i.e., from *v* to (*i, c*)) are performed while looping over the vectorized coordinates, i.e., the client can decrypt and decode the annotation values as they are received from the annotation server in a streaming fashion.

For SNVs, the client performs the following steps:

1. Receive and decrypt data
2. For each non-zero value on the annotated vector at position *v*,
3. Translate *v*: Compute *i*^(*T*)^(*v*) and *c*^(*T*)^(*v*)
4. Extract the 38-bit impact bitmap from the annotated value
5. For every bit that is set to 1, use the lookup table and concatenate the variant annotation.

### Secure Small Deletion Variant Annotation

Annotation of deletion variants needs to account for the variable lengths of the variants. The server can provide all the vectorized annotation signals for deletions that are shorter than a certain value, e.g., 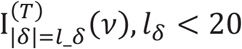. Similarly, the researcher can generate the encrypted variant loci vectors for deletions of each length. After this, the annotation can be performed as it is performed for SNVs annotations by first secure multiplication of the variant loci vector and annotation vector for each deletion length, then decoding the 38-bit annotation impact bitmap for each non-zero entry in the product vector for each length.

SVAT also implements an alternative approach for the annotation of deletions. SVAT uses the 1-base pair deletion impact signal to track the impact of consecutively deleted nucleotides: For a deletion of nucleotides [*a, b*] (*a < b*), SVAT traces the nucleotides and merges the impacts of consecutive 1-base pair deletions, at every nucleotide *v* in [*a, b*], i.e., 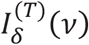. To merge the 1-base pair deletion annotations and to generate the impact of a deletion at [*a, b*], SVAT makes use of the coding frame information and the neighborhood sequence.

#### Coding Frame Impact

SVAT traces the impact values of all nucleotides between [*a, b*], and counts the number of coding nucleotides that are affected by the deletion. If the number of coding nucleotides is a multiple of 3, the deletion is marked as an in-frame deletion. Otherwise, the deletion is marked as a frameshift deletion.

#### Start/Stop Loss

If any of the nucleotides in [*a, b*] have an impact on the start or stop codons, the deletion is marked with a start/stop loss.

#### Splice Donor/Acceptor Loss

If any of the nucleotides have an impact on the splice donor/acceptor sites, the deletion is marked with splice donor/acceptor.

#### Stop Gain/Retain

SVAT utilizes the neighborhood and the coding frame to identify the coding frame around the deletion breakpoint. This way, SVAT computes the new codon (using a lookup table) that is introduced by the deletion. If the new codon is one of the three stop codons, i.e., “TAG”, “TAA”, “TGA”, SVAT assigns the stop gain impact. If the deletion impacts a stop codon and generates a new one around the breakpoint, “Stop Retained” impact is added to the aggregate impact value of the variant.

### Secure Small Insertion Variant Annotation

Insertions are short sequences that are inserted with respect to the reference genome. Unlike a deletion, an insertion occurs at a single location and is described by the sequence that is inserted at the location. The annotation of insertions is carried out the same way as deletions by first identifying the position of the insertion and then translating the coding nucleotides that are inserted to assign the impact.

#### Coding Frame Impact

SVAT traces the number of coding nucleotides that are inserted into a CDS and assigns in-frame or frameshift annotation.

#### Start/Stop Loss/Gain/Retainment

For any insert that overlaps with a start or a stop codon, the neighborhood nucleotide and the coding frame is used to identify the inserted codons and set whether a stop/start codon is lost/gained or retained.

The scenario of secure insertion annotation is the same as secure deletion annotation, where 1-base pair insertion annotation vector 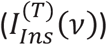 is multiplied by the encrypted 1-base pair insertion mutation vector 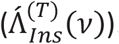. The multiplied encrypted vector is sent back to the researcher, which is decrypted and processed.

### Genotype Aggregation

The genotype aggregation aims as computing the frequencies of the mutations by aggregating over many samples:

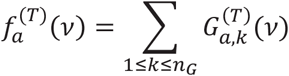

where *n*_*G*_ denotes the number of individuals in the genotype matrix, 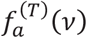 denotes the frequency of the allele *a* for the variant at vector coordinate *v* (indexed on the target regions *T*), and 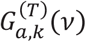 indicates the genotype of the variant for individual 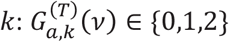, where the entry indicates the number of alternate alleles equal to *a*. The genotypes can also be encoded with 2-level encoding[39] to indicate the existence of the allele at the location: 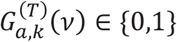.

While aggregating the genotype matrix, it is necessary to ensure the confidentiality of the genotype matrix 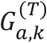 from the untrusted aggregation server, which performs the computationally heavy task of secure aggregation. To compute the above summation securely, the genotype matrix is encrypted, denoted by 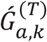 and the summation is evaluated using HE-based secure summation.

As practical homomorphic encryption systems support computation on encrypted vectors, we encrypt the whole genotype matrix 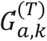 by taking entries in row-major order and generating plaintext vectors. In other words, each row vector is divided into sub-vectors of length *l*, each of which is encrypted using the public key. This vectorization makes it possible to use SIMD instructions that operate the aggregation on vectors (i.e., with support for entry-wise addition). To be specific, suppose that for each *k* 1,2, …, *n*_*G*_, a plaintext vector 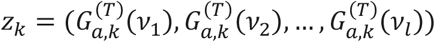 is encrypted into a single ciphertext *ct*_*k*_. Then we perform aggregation over ciphertexts: 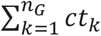. By the homomorphic property, we have

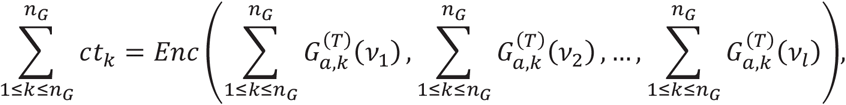

which is an encryption of the frequency vector 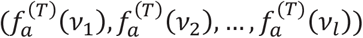. The ciphertext packing technique enables us to encrypt *l* different messages into a single ciphertext and perform *l* operations at a time over encryption.

Another important aspect of aggregation is to accommodate genotype matrices from multiple databases so that many samples can be aggregated together. The server needs to be able to manage multiple datasets that are encrypted with different keys. SVAT implements a proxy re-encryption protocol to convert the genotype matrices into the same key and perform the aggregation using this common key.

#### Proxy re-encryption protocol

The proxy re-encryption aims at converting the genotype matrices (or any other type of data) to the same encryption key[65]. For this, a trusted entity is required who will perform the key management who holds the private keys necessary to generate the re-encryption keys. This is a reasonable assumption since the sensitive datasets are generally deployed and protected by entities such as NIH (e.g., dbGAP).

We assume that there are *M* genotype matrices (on same vectorized coordinates system, i.e., common sorted target regions *T*), such that *m*^*th*^ matrix is encrypted with the public key denoted by 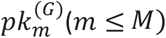. The corresponding private keys, 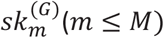, are stored by the trusted entity such as NIH, and *m th* matrix can be decryptable only with the private key *sk m*(*G*). The data is stored at the untrusted server in an encrypted format. When a researcher asks for the aggregation service, the researcher provides the public key, *pk*^(*R*)^ (Researcher public-key) and a corresponding private key *sk*^(*R*)^. It is necessary to re-encrypt the encrypted genotype matrices 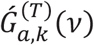 so that they can be decrypted with the private key *sk*^(*R*)^. The researcher sends the public and private keys to NIH, which generates the re-encryption key for the data, i.e. *swk* 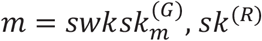 for each *m* ≤ *M* where *swk m* denotes the re-encryption key. The aggregation server receives *swk m* for all genotype matrices and re-encrypts so that they can be decrypted by the same key of the user *sk*^(*R*)^.

Traditional homomorphic encryption systems can convert a ciphertext decryptable with a secret key *sk*_1_ to a new ciphertext with a secret key *sk*_2_, which is called the *key-switching* operation. To be specific, if the switching key *swk* = *swk*(*sk*_1_, *sk*_2_) is generated, then the key-switching operation takes as input a ciphertext *ct* with the secret key *sk*_1_ and perform the key-switching operation

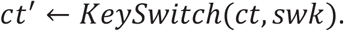

Then the resulting ciphertext *ct*^′^ is decryptable with the secret key *sk*_2_. In our case, the secret keys are set as 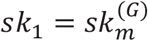 and *sk*_2_ = *sk*^(*R*)^, and the aggregation server performs the key-switching operations on the encrypted genotype matrices using the public switching key *swk*_*m*_ without knowing any information about the secret keys. The aggregation service then securely aggregates the frequency counts at every position on the *T*. The resulting frequency array, which is encrypted with the researcher’s public key, 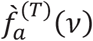, is sent to the researcher who can decrypt the frequency array and obtain frequencies.

### SNV Aggregations

The secure aggregation of the SNVs is straightforward as they are located at one position on the vectors. In other words, the SNV frequency aggregation can be computed simply by marginalizing at every location.

### Indel Aggregations

Unlike SNVs, the deletions must be tracked in each sample and aggregated. As with annotation task, we make use of the 1-base pair deletions to build the aggregation of deletion variants that span longer than 1bp. Given a position *v*, and the 1-base pair deletion genotype matrix, 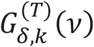; the indel of length *l*_*δ*_ is aggregated by tracing the deleted nucleotides:

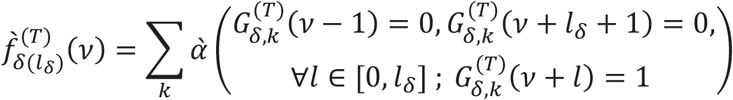

where 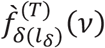 indicates the frequency of the *l*_*δ*_-deletion at vector position *v*, and *α* (·) is an indicator function, i.e., it returns 1 if the arguments are true and 0 otherwise. In the formula above, the summation aggregates the individuals for which, the 1-base pair deletion genotype matrix contains a deletion state for all positions in {*v, v* + 1, *v* + 2, …, *v* + *l*_*δ*_}. In addition, the deletion state is unset (i.e., 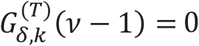) for the individual *k* at positions {*v* − 1, *v* + *l*_*δ*_ + 1}. This ensures that the deletion is set exactly over the deleted interval we would like to aggregation, i.e., [*v, v* + *l*_*δ*_] and also the positions right out of the deleted interval are set to undeleted. It should be noted that for any aggregation of deletions, the aggregation can be performed for multiple deletion lengths (i.e., *l*_*δ*_ < *l*_*max*_) and stored at the aggregation server.

The aggregation of insertions requires explicit matching of the inserted nucleotides and requires enumeration of all possible insertions. SVAT currently does not explicitly support aggregation of short insertion variants. However, the position at which the insertion happens can be aggregated (just by simple aggregation as for SNVs) to compute the frequency of insertion at each position.

## List of Abbreviations

We list the variables used in the text for easy accession to the readers.

*i*: A sorted target region index.
*c*: A genomic coordinate.
*pos*: A genomic coordinate.
*v*: Index of the vectorized coordinates
*T*: Sorted target regions list
*s*_*i*_: Genomic position for the start of *i*^*th*^ target region.
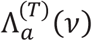: Variant loci vector of length *l*_*T*_ for allele *a, a ϵ* {*A, C, G, T, δ, l*} where *8* and *L* indicate 1-base pair deletion and 1-base pair insertion.
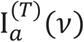: Variant impact vector of length *l*_*T*_.
*e*_*i*_: Genomic position for the end of *i*^*th*^ target region.
*l*_*ext*_: Extension length by which each target has been extended at start and end.
*v*_*<k*_: The total coverage of the target regions to the left of *k*^*th*^ sorted target region.
*i*^(*T*)^(*v*): Index of the target region whose coordinates overlap with the vectorized coordinate *v*.
*c*^(*T*)^(*v*): Genomic coordinates for the vectorized coordinate *v*.
*N*^(*T*)^(*pos*): The set of vectorized coordinates for which the genomic coordinate is *pos*.

## Acknowledgements

We acknowledge and thank Luyao Chen for providing technical support to set up the computational environment for performing benchmarking experiments.

## Authors’ information

Not applicable.

## SVAT: Secure Variant Annotation and Aggregation Tool

